# Rare Epithelial Sentinel Cells Establish Functional Domains of the Airway

**DOI:** 10.64898/2026.11.20.768192

**Authors:** Jiajie Xu, Matthew Ho, Regina V. Oliveira, Ella Jacobs, Jennifer L. Reedy, Kirstine Nolling Jensen, Haitian Zhao, Philippe Marambaud, Michael K. Mansour, Raul Mostoslavsky, Jatin M. Vyas, Jayaraj Rajagopal

## Abstract

Organs exposed to the environment must sense threats in order to evoke protective physiologic responses. Recent single-cell sequencing studies of the airway epithelium have led to the identification of a suite of rare epithelial cells, many of which are unified by their ability to sense stimuli^1,2^. Herein, we explore how organized networks of epithelial sentinel cells transduce distinct stimuli to alter tissue function by employing a combination of cell type-specific calcium indicators, optogenetics, chemogenetics, cell ablation, and live ACh and ATP sensors. We define a new diversity of functional sentinel cells including 3 new subtypes of chemosensory tuft cells. We further show that solitary neuroendocrine cells, like their tuft cell counterparts, generate stimulus-induced waves of calcium activation. These waves originate from a central sentinel cell and expand outwards to encompass a circular domain of surrounding epithelial cells. We demonstrate that basolateral secretion and diffusion of neurotransmitters cause the activation of the surrounding epithelial effector cells including basal stem cells which were not previously known to be participants in sentinel cell-induced calcium waves. We show that these diffusion-limited domains (DLDs) are generated by specific patterns of ACh and ATP secretion characteristic of specific sentinel cell populations. The speed, intensity, and domain size of DLDs are dictated by the degree of stimulus-triggered neurotransmitter secretion. Thus, DLDs represent a unified mechanism for specifying functional domains within the airway epithelium.

Triggered sentinel cells generate surrounding circular domains of increased mucociliary transport and enhance stem cell replication. Insulated sentinel cell networks are tuned to distinct stimuli. Sentinel cells are distributed in sufficient numbers to ensure comprehensive epithelial coverage. However, the amplitude and number of DLDs scale with stimulus intensity, permitting both local and global physiologic activation of the epithelium. Additionally, the distribution of sentinel cells can vary with organ anatomy, resulting in compartmentalized physiologic responses. Furthermore, cytokines modulate DLD responses. IL-1β amplifies ciliary action by generating larger and more intense DLDs, while IFN-γ reduces the density of DLDs by directly depleting sentinel cells. Finally, we show that a fungal metabolite triggers DLDs. In aggregate, we reveal a sophisticated collective of diverse sentinel cells that activate surrounding circular domains of airway epithelium to generate protective responses to environmental threats.

---

The mammalian lung is composed of two major compartments: the airways that conduct gases to the more distal alveoli that perform gas exchange. Thus, inhaled threats are first sensed in the airway. We recently identified a new hierarchy of airway epithelial cells, including historically recognized solitary neuroendocrine cells (NE) and two subtypes of tuft cells of unclear significance^1,2^. NE cells and tuft cells have previously been shown to participate in the regulation of mucociliary transport, airway inflammation, and respiratory reflexes^3–16^. NE cells and tuft cells are also known to be responsive to environmental stimuli. Airway tuft cells are responsive to bitter taste receptor ligands^4–8,12^, bacterial quorum-sensing molecules^4,5,17^, formyl peptides^3^, and succinate^11^, while airway solitary NE cells sense mechanical stimuli, water, and acid, as well as succinate^9,10^.

Herein, we employ live calcium imaging to assess the aggregate behavior of ensembles of airway epithelial cells because NE cell and tuft cell activation have both been associated with calcium transients^3,9^. More than three decades ago, mechanical stimulation of the airway epithelium was shown to result in a wave of increasing Ca^2+^ that spread, cell by cell, from the stimulated cell to neighboring cells^18^. Recently, airway tuft cells have also been shown to initiate ICWs that activate neighboring ciliated and secretory cells^11,12^. Indeed, the phenomenon of intercellular calcium waves (ICWs) has been widely documented in many organs^19^, but their functions are largely obscure.

We deploy live imaging coupled to optogenetics, chemogenetics, cell type-specific ablation, and live sensors of neurotransmission to dissect the molecular and cellular mechanisms governing how environmental stimuli engender tissue-wide physiologic responses. First, we demonstrate that solitary NE cells engender waves of calcium activation mirroring the behavior of their tuft cell counterparts. We hypothesize that the functional units of the airway epithelium are organized as circular domains of effector cells surrounding a distributed network of diverse sentinel cells. We show that tuft and NE cell networks, triggered by distinct stimuli, are functionally insulated from one another. Furthermore, sentinel cell networks are not uniformly distributed throughout the airway, but are instead patterned by the varying anatomic distribution of sentinel cells. Adding to the complexity, we define a diversity of functional tuft cells that underpin distinct networks of effector cells responsive to distinct stimuli. In contrast to the prevailing gap junction model for the generation of tuft cell-initiated calcium waves^11,12^, we show that tuft cells and NE cells both employ varying configurations of basolateral neurotransmitter secretion to generate diffusion-limited domains (DLDs) of calcium activation. Furthermore, we demonstrate for the first time that stem cells, like their neighboring ciliated and secretory cells, are functionally activated by triggered sentinel cells. Thus, DLDs represent distinct domains of airway epithelium in which the two cardinal functions of the airway epithelium, mucociliary transport and epithelial self-renewal, are physiologically coupled. Furthermore, disease-relevant cytokines can enhance or limit DLDs and their corresponding physiologic outputs. Finally, we demonstrate that a clinically relevant respiratory pathogen is detected by a single sentinel cell receptor whose activation in turn triggers DLDs.

## Unique stimulus-induced intercellular calcium waves (ICWs) originate from both tuft and NE sentinel cells and are insulated from one another

Previous studies of airway epithelial calcium signaling mostly relied on calcium indicator dyes that did not resolve the behavior of distinct cell types^11,12,18,20^. Therefore, we employed the genetically encoded calcium indicator GCaMP^21,22^, under the control of pan-epithelial and cell-type-specific drivers to monitor calcium dynamics. We established an air-liquid interface (ALI) culture system that permits the use of an additional red calcium indicator XCaMPR^23^, the optogenetic actuator mOPN4^24,25^, and live sensors for neurotransmitter acetylcholine (ACh)^26^ or ATP^27^ (Extended Data Fig. 1a). This integrated toolkit enabled us to dissect the molecular and cellular mechanisms of the airway physiologic response to environmental stimuli.

We first confirmed that succinate induced robust ICWs as previously reported^11^ (Fig. 1a, Extended Data Fig. 2, and Supplementary Video 1). We then determined the concentration-response curve with an EC50 of 116 µM using our *in vitro* platform (Extended Data Fig. 3a). To establish physiologic relevance, we confirmed our findings in an airway explant model that we developed (Extended Data Fig. 1b, 3c). These ICWs originated from discrete foci distributed throughout the epithelium and propagated to eventually cover the entire tissue (Extended Data Fig. 3c). Thus, the pattern of tuft cell distribution dictates a pattern of overlapping ICWs that encompasses the entirety of the airway lining. To verify the identity of the initiating cells, we performed live imaging followed by fixation and immunostaining with cell-type-specific markers. By registering the fixed images to the live recordings, we identified a tuft cell at the center of every succinate-induced ICW (Fig. 1a).

**Figure 1.**
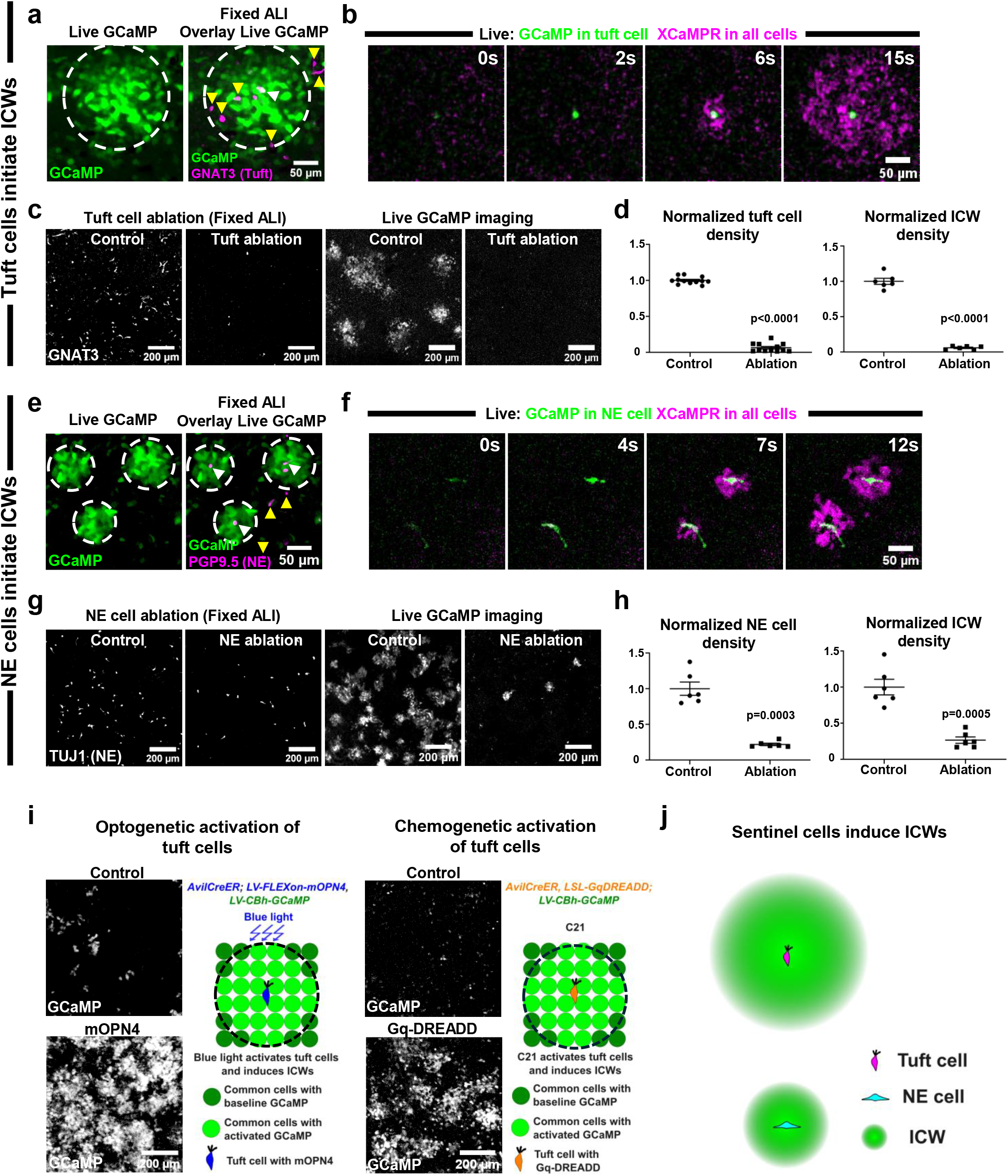
Stimulus-induced intercellular calcium waves (ICWs) originate from both tuft and NE sentinel cells. (**a**) Succinate application induces an ICW (ΔF/F GCaMP signal) in an ALI culture. Post-imaging immunostaining for the tuft cell marker GNAT3 identifies a tuft cell (white arrowhead) at the center of the ICW. Yellow arrowheads indicate non-ICW-inducing tuft cells. The white dashed circle denotes the ICW domain. (**b**) Dual-color live imaging (GCaMP in tuft cells; XCaMPR throughout the epithelium) reveals sequential tuft cell activation (green) followed by radial ICW propagation (magenta) in response to succinate. (**c, d**) Genetic ablation of tuft cells (*AvilCreER; LSL-DTA*) efficiently depletes GNAT3+ tuft cells and abrogates succinate-induced ICWs. Welch’s t-test in (d). (**e**) High K^+^ application induces three ICWs. Post-imaging staining for the NE cell marker PGP9.5 identifies a NE cell (white arrowhead) at the center of each ICW. Yellow arrowheads indicate non-ICW-inducing NE cells. White dashed circles denote ICW domains. (**f**) Dual-color live imaging (GCaMP in NE cells; XCaMPR throughout the epithelium) shows sequential NE cell activation (green) followed by radial ICW propagation (magenta) in response to high K+. (**g, h**) Genetic ablation of NE cells (*Ascl1CreER; LSL-DTA*) results in a 78% reduction of TUJ1+ NE cells and a corresponding decrease in high K+-induced ICWs. Welch’s t-test in (h). (**i**) Optogenetic and chemogenetic activation of tuft cells is sufficient to induce robust ICWs. The black dashed circles in the schematics denote the ICW domains. (**j**) Schematic model summarizing the role of tuft and NE cells as initiators of ICWs.

In order to isolate the initial activation of a sentinel cell from the subsequent propagation of the calcium wave without the need for fixation and post-hoc registration, we developed a live dual-color calcium imaging regime. Using an *AvilCreER*^28^ driver line, we expressed green GCaMP in tuft cells and additionally employed lentiviral transduction to express the red calcium indicator XCaMPR^23^ throughout the epithelium. We were able to visualize the sequential activation of individual tuft cells preceding the radial propagation of the ICW (Fig. 1b). We also found that only a minority of tuft cells initiated ICWs. Indeed, tuft cells exhibited three distinct response patterns to succinate: no activation; activation without ICW initiation; and activation with ICW initiation (Extended Data Fig. 3b).

We then sought to confirm the necessity of tuft cells in generating ICWs using cell type-specific genetic ablation with Diphtheria Toxin A (DTA)^29^. This ablation completely abrogated succinate-induced ICWs (Fig. 1c, d). We bypassed the need for the use of a natural ligand that may have broad effects on the epithelium using two orthogonal methods: chemogenetic activation with Gq-DREADD^30^ and optogenetic activation with mOPN4^24,25^. Both approaches robustly induced ICWs in ALI cultures (Fig. 1i, Extended Data Fig. 4, and Supplementary Video 2, 3), demonstrating that a calcium spike is sufficient to bypass the need for a specific ligand to induce an ICW.

We next investigated whether solitary NE cells, a second rare chemosensory cell type, initiate ICWs. NE cells have recently been shown to fire action potentials in response to a high K^+^ bath^9^. We show that a high K^+^ bath induced ICWs in both ALI cultures (Fig. 1e and Supplementary Video 4) and explants (Extended Data Fig. 3d). A NE cell was located at the center of every high K^+^ induced ICW (Fig. 1e). Using our dual-color calcium imaging strategy with GCaMP expressed specifically in NE cells (*Ascl1CreER*^31^) and XCaMPR in all cells, we confirmed that NE cell activation consistently preceded ICW propagation (Fig. 1f). Furthermore, genetic ablation of NE cells (78% efficiency) resulted in a corresponding reduction in high K^+^ induced ICWs (Fig. 1g, h). In contrast to tuft cells, a majority of NE cells initiated ICWs. NE cell-initiated ICWs were consistently smaller and propagated more slowly than those initiated by tuft cells (Fig. 2a).

**Figure 2.**
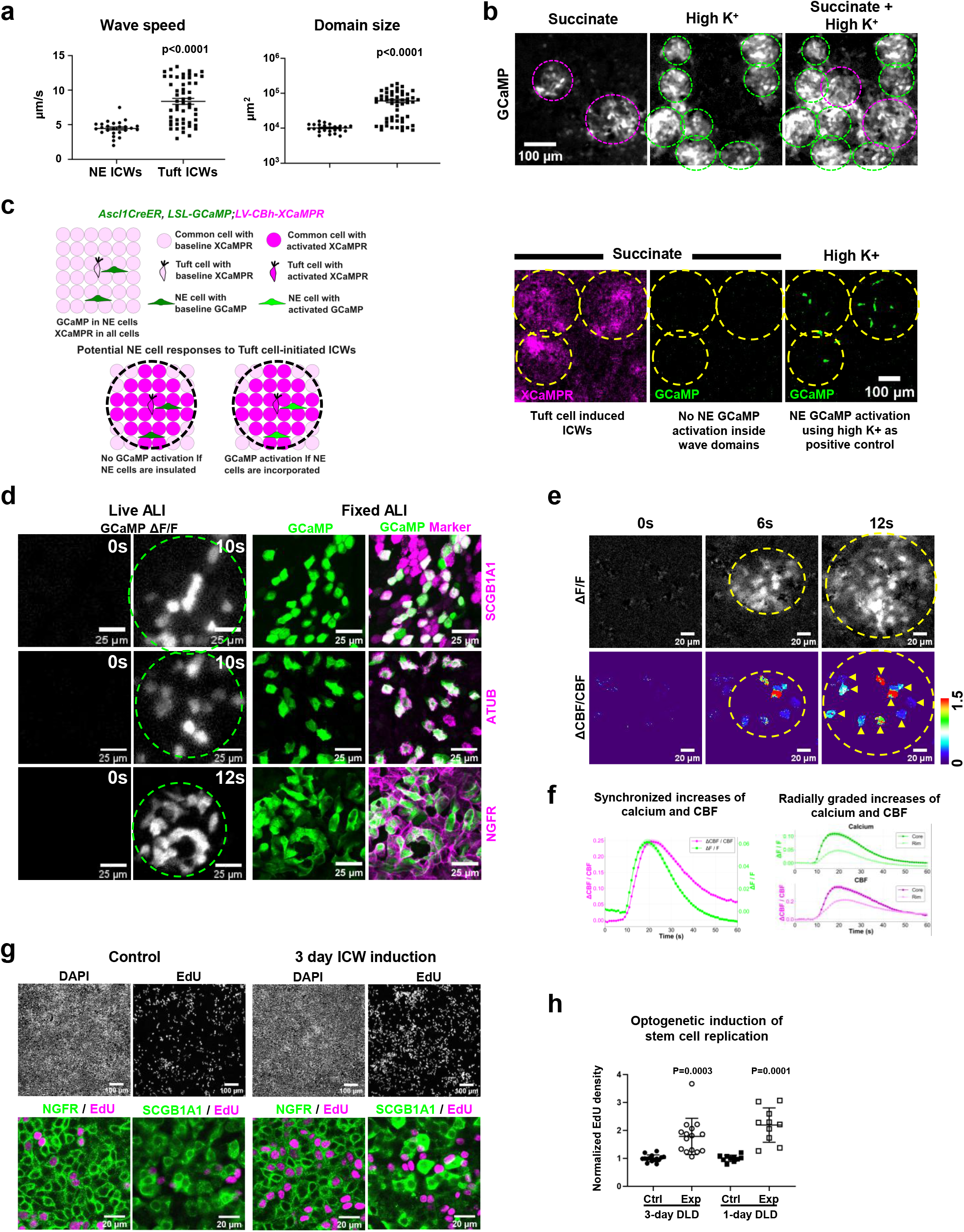
Stimulus-triggered tuft cell and NE cells generate distinct functional domains of the airway epithelium and increase stem cell replication. (**a**) Comparison of wave speed and domain size for ICWs initiated by NE cells versus tuft cells (Welch’s t-test). (**b**) Co-application of high K^+^ and succinate generates both NE cell- and tuft cell-initiated ICWs within the ALI culture. The magenta dashed circles denote tuft cell-initiated ICW domains whereas the green dashed circles denote NE cell-initiated ICW domains. (**c**) NE cells located within tuft cell-initiated ICW domains do not exhibit calcium spikes, despite being responsive to direct high K^+^ depolarization, indicating that NE cells are insulated from tuft cell-initiated ICWs. The black dashed circles in the schematics and the green dashed circles in live imaging denote tuft cell-initiated ICW domains. (**d**) Calcium imaging with cell type-specific GCaMP expression reveals that high K^+^-induced ICWs activate secretory, ciliated, and basal stem cells, indicating that all three common cell types participate in NE cell-initiated ICWs. Identities of GCaMP expressing cells are confirmed by staining of common cell markers: SCGB1A1 for secretory cells, Acetylated Tubulin (ATUB) for ciliated cells, and NGFR for basal stem cells. The green dashed circles denote NE cell-initiated ICW domains. Time stamp in seconds shown at top right corner. **(e)** High-speed live imaging of NE cell-initiated ICWs and ciliary beat frequency (CBF). Top row: calcium activation visualized as ΔF/F of the GCaMP signal within the ICW domain (yellow dashed circle). Bottom row: ΔCBF/CBF is coupled to the ciliated cells calcium activity. Time stamps in seconds shown at top. **(f)** Quantitative traces of calcium (ΔF/F, green) and CBF (ΔCBF/CBF, magenta) within NE cell functional domains. Top: aggregate response across the entire domain. Bottom: separation of responses into domain core (cells within half the wave radius) and domain rim (cells beyond half the wave radius) reveals larger amplitudes in the core for both calcium and CBF. **(g)** Three-day optogenetic induction in *AvilCreER; LV-FLEXon-mOPN4* ALI cultures drives EdU incorporation only in NGFR⁺ basal stem cells, but not in SCGB1A1⁺ secretory progenitor cells. **(h)** Normalized EdU⁺ cell density in control vs. optogenetically activated cultures showing significant proliferative responses at both day 1 and day 3 (Welch’s t-test).

We next sought to explore potential functional interactions between tuft cell-initiated and NE cell-initiated ICWs. Co-application of high K^+^ and succinate generated both types of ICWs within the same culture, demonstrating their coexistence (Fig. 2b). To test if there is direct communication between tuft cells and NE cells through calcium signaling, we expressed GCaMP in both tuft cells and NE cells and asked if activation of one cell type leads to activation of the other. We found that tuft cells and NE cells are insulated from each other (Extended Data Fig. 5a). To more rigorously assess whether NE cells or tuft cells could be activated by neighboring ICWs originating from the opposing cell type, we first imaged GCaMP-expressing NE cells in the setting of succinate-induced tuft ICWs visualized by XCaMPR. NE cells located within the domains of tuft cell-initiated ICWs did not display calcium spikes, although they did respond to direct depolarization with high K^+^ (Fig. 2c). Similarly, the majority of tuft cells located within NE cell-initiated ICWs did not display calcium spikes (Extended Data Fig. 5b). Thus, these two sentinel cells are insulated from activation by ICWs originating from the opposing sentinel cell type.

To precisely determine which common cell types are responsive to sentinel cell activation, we generated ALI cultures with GCaMP expression restricted to individual cell lineages: secretory (*Scgb1a1CreER*^32^), ciliated (*Foxj1Cre*^33^), or basal stem (*P63CreER*^34^) cells. Consistent with previous observations, secretory and ciliated cells were recruited into ICWs^11^. Surprisingly, we now demonstrate that basal stem cells also participate in the ICWs (Fig. 2d, Extended Data Fig. 6). Thus, every common cell type participates in ICWs. The inclusion of basal stem cells suggests that sensed stimuli may prime subsequent regenerative events.

## Sentinel cells generate distinct functional domains of the airway epithelium and activate stem cell replication

The airway epithelium has two defining functions: it clears inhaled material through coordinated ciliary beating, and it regenerates lost cells through stem cell activity. Recent work has shown that succinate activation of tracheal tuft cells leads to increased ciliary beat frequency (CBF)^11^. We sought to determine whether local domains of calcium activation result in spatially restricted zonal physiologic responses. To do so, we measured CBF within individual ICWs using high-speed live imaging. Tuft cell-initiated ICWs were evoked by succinate, and produced a robust increase in CBF (Extended Data Fig. 7a). Both calcium (ΔF/F) and CBF (ΔCBF/CBF) were synchronized. Indeed, markedly larger amplitudes of CBF were elicited in the core of the ICWs (cells within half the wave radius) when compared to the rim (cells beyond half the wave radius) (Extended Data Fig. 7b). Furthermore, we show that NE cell-initiated ICWs evoked by high K⁺ produced the same pattern of local physiologic response (Fig. 2e, f). Thus, physiologic responses of the airway epithelium occur in zonal domains, irrespective of the initiating sentinel cell type and the specific sensed stimulus.

Unlike secretory and ciliated cells, stem cells were not previously associated with sentinel cell-induced ICWs. Optogenetic induction of ICWs over 3 days resulted in a significant increase in EdU incorporation in NGFR⁺ basal stem cells, but had no effect on the SCGB1A1⁺ secretory cells, which are also known to possess replicative capacity^32,35^ (Extended Data Fig. 8a, Fig. 2g, h). Remarkably, optogenetic stimulation was sufficient to drive stem cell replication within 1 day (Fig. 2h, Extended Data Fig. 8b).

## Acetylcholine (ACh) diffusion delimits tuft cell-initiated ICW domain size

Recent work proposed a sequential model for the propagation of tuft cell-initiated calcium waves in the trachea, in which paracrine ACh from tuft cells first activates immediately adjacent cells, and the wave is then carried to distant cells through gap junction communication^11^. In this model, the domain size of the wave reflects the reach of gap junction-mediated propagation, with ACh acting only on the immediate neighbors of the initiating tuft cells^11^. To rigorously evaluate the contribution of each step to the spatial properties of tuft cell-initiated ICWs, we systematically tested both the diffusion range of ACh itself and the requirement for gap junction-mediated propagation. We employed a newly available live ACh sensor ^26^ in ALI culture via lentiviral transduction to visualize the diffusion of ACh. Succinate-induced waves of ACh sensor activation originated from discrete foci and extended over large domains, encompassing more than ten cell diameters (Extended Data Fig. 9a). To precisely correlate ACh activity with increases in calcium, we developed a dual-sensor system, allowing for simultaneous imaging of the green ACh sensor and the red calcium indicator XCaMPR. This revealed that the wavefront of ACh activity consistently preceded the wavefront of calcium elevation by several seconds (Fig. 3a and Supplementary Video 5). This temporal relationship is consistent with a model where ACh diffusion and subsequent receptor activation directly drive the ICW. Indeed, the domain size of ACh sensor activity directly corresponds to the size of the calcium waves (Fig. 3a).

**Figure 3.**
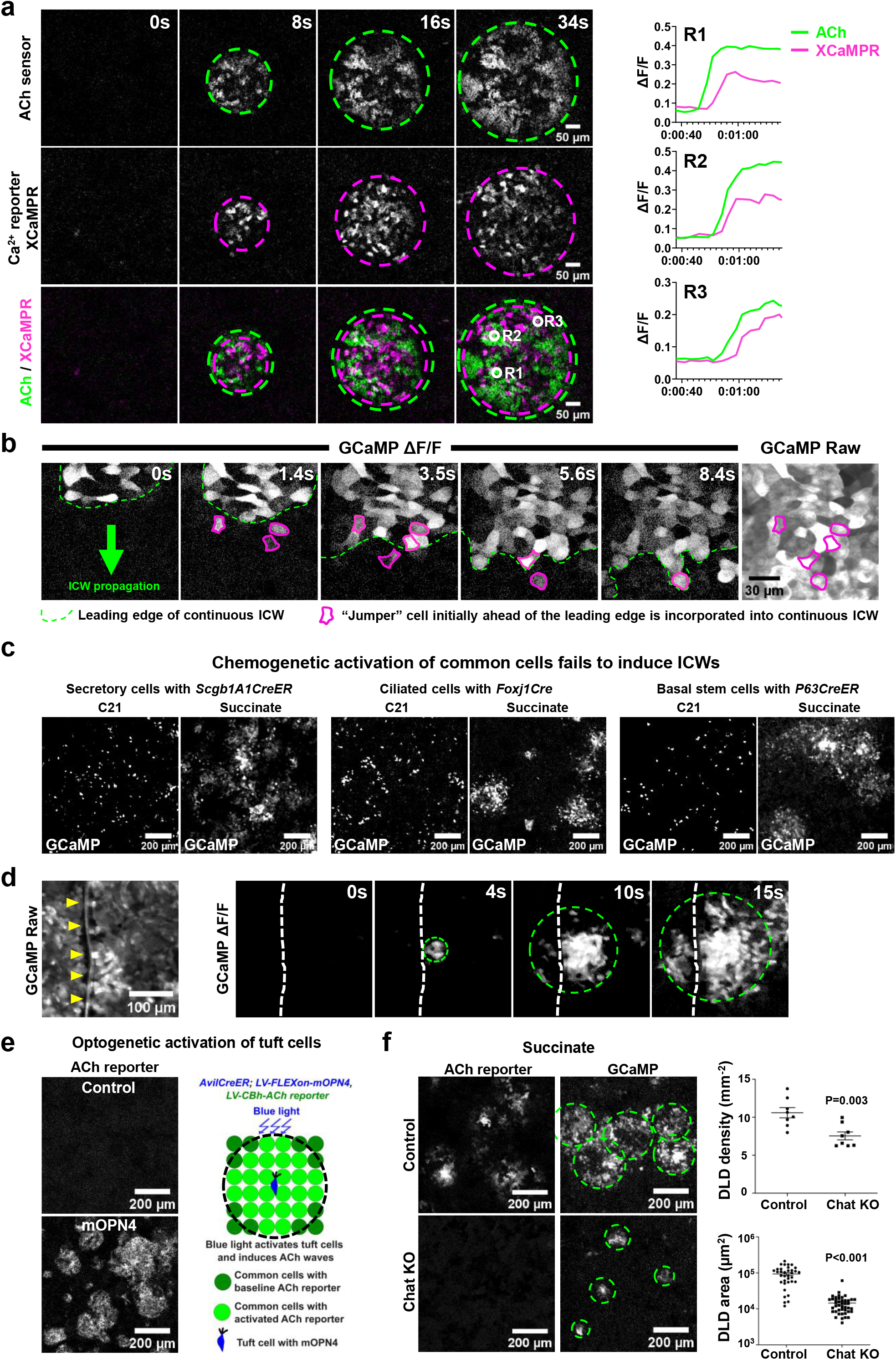
Acetylcholine (ACh) diffusion delimits tuft cell-initiated ICW domain size. (**a**) A dual ACh/Ca²⁺ sensor reveals that the Ca²⁺ wave (magenta dashed circles) slightly lags behind the diffusion of ACh (green dashed circles) following succinate application. Traces from three regions (R1-R3) show a several-second delay between ACh and Ca²⁺ increases. (**b**) “Jumper” cells (outlined in magenta) exhibit calcium spikes ahead of the leading edge of the continuous ICW following succinate application. Raw GCaMP fluorescence shown on the right confirms GCaMP expression in cells surrounding jumper cells. The green arrow indicates the direction of ICW propagation. (**c**) Sparse chemogenetic activation of common epithelial cells using the DREADD ligand C21 triggers isolated calcium spikes but fails to initiate ICWs, whereas subsequent succinate application induces robust ICWs in the same tissue. (**d**) ICWs propagate across a physical cut created by a tungsten needle, indicating that direct cellular continuity is not required for wave transmission. Yellow arrowheads and white dashed lines indicate the cut site. (**e**) Optogenetic activation of tuft cells is sufficient to initiate ACh waves. (**f**) Residual ICWs persist after the genetic ablation of ACh signaling. In *Chat* KO cultures, ACh sensor signal is absent (left), and ICWs are reduced in number and domain size (right; Welch’s t-test).

Our high-resolution live imaging of ICW propagation revealed another key finding that supports the diffusion model: the presence of insulated “jumper” cells with calcium spikes ahead of the leading edge of the continuous wave. The discontinuity implies that they could not have been activated by gap junctions that operate through direct physical contact (Fig. 3b). Importantly, GCaMP expression in neighboring cells was confirmed to rule out the possibility that the “jump” is an artifact of patchy GCaMP expression. This phenomenon is fundamentally incompatible with a sequential gap junction propagation mechanism, in which a cell can only be activated after a directly adjacent cell has been activated. To further distinguish between gap junction-mediated and diffusion-mediated wave mechanics, we sparsely expressed the excitatory Gq-DREADD in each of the common cell types (secretory, ciliated, and basal stem cells). In contrast to succinate application, the application of the DREADD agonist compound 21 (C21) induced calcium spikes only within the targeted cells, with no propagation of calcium wave to neighbors as expected in the gap junction model (Fig. 3c, Extended Data Fig. 10). Finally, we reasoned that gap junction coupling would be prevented by a physical separation of the epithelium. We used a tungsten needle to create an incision in ALI cultures. Strikingly, ICWs propagated seamlessly across the cut edge (Fig. 3d and Supplementary Video 6), consistent with a simple diffusion model. Using an orthogonal approach, we show that optogenetic activation of tuft cells elicits robust ACh-mediated waves (Fig. 3e). These results strongly favor a simple model in which ACh diffusion dictates the domain size of an ICW. Thus, airway “ICWs”, previously thought to represent gap junction-mediated “waves” of calcium activation, are instead domains that are defined by simple neurotransmitter diffusion, hereafter referred to as diffusion-limited domains (DLDs).

## Diverse sentinel tuft and NE cells employ varying combinations of basolateral ACh and ATP secretion to pattern diverse diffusion-limited domains (DLDs)

To further explore the ACh diffusion model, we generated conditional *Choline acetyltransferase knockout* (*Chat KO*) in tuft cells to eliminate ACh synthesis using the Chat^flox/flox^ allele ^36^ (Fig. 3f). Surprisingly, *Chat KO* resulted in the elimination of only a quarter of succinate-induced DLDs (Fig. 3f). The remaining three-quarters of DLDs were markedly smaller in domain size. This puzzling result led us to hypothesize that a majority of DLDs are propagated by an additional, ACh-independent signaling molecule that diffuses over a smaller domain, a possibility we explore next.

Taste cells share a common transduction machinery with airway tuft cells and secrete ATP through the CALHM1 channel to activate sensory nerves^37–39^. More recently, tuft cells in the hypopharynx and larynx have been shown to secrete ATP to communicate with vagal sensory nerves^8^. Furthermore, previous studies have shown that ATP can induce waves of calcium activation in airway epithelium^40^. Therefore, we hypothesized that ATP secretion was responsible for the aforementioned small residual DLDs. To test this hypothesis, we employed a recently available live ATP sensor^27^, and demonstrated that succinate triggered ATP waves (Extended Data Fig. 9b). These ATP waves were significantly smaller than the ACh waves. Furthermore, optogenetic activation of tuft cells alone was sufficient to elicit ATP waves (Fig. 4c).

**Figure 4.**
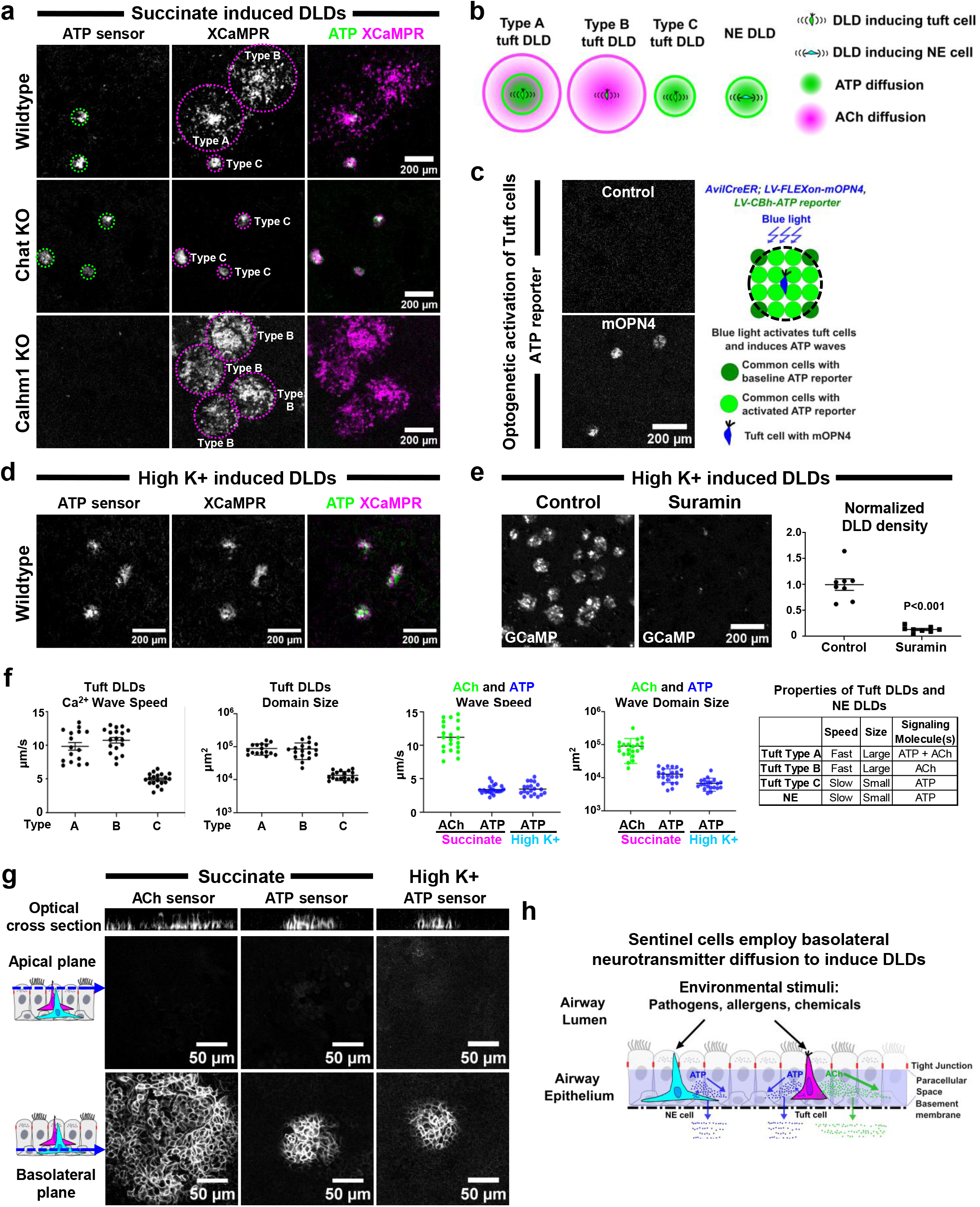
Distinct sentinel cells employ distinct neurotransmitter release mechanisms to pattern diverse diffusion-limited domains (DLDs). **(a)** Characterization of succinate-induced DLDs using an ATP/Ca²⁺ dual sensor reveals three distinct types. Type A: A small ATP wave (green) surrounded by a larger Ca²⁺ wave (magenta). Type B: A large Ca²⁺ wave with no detectable ATP wave. Type C: A small Ca²⁺ wave perfectly co-localized with an ATP wave. Genetic ablation of *Chat* eliminates Type A and B DLDs, leaving only Type C, whereas *Calhm1* KO abolishes Type A and C, leaving only Type B. **(b)** Schematic model summarizing different types of DLDs. **(c)** Optogenetic activation of tuft cells is sufficient to induce ATP waves. **(d)** ATP/Ca²⁺ dual sensor imaging shows that high K⁺-induced NE cell DLDs are characterized by co-localizing ATP (green) and Ca²⁺ (magenta) waves. **(e)** Pharmacological blockade of P2 purinergic receptors with suramin abrogates high K⁺-induced ICWs, demonstrating that ATP signaling is necessary for NE DLD propagation (Welch’s t-test). **(f)** Quantitative summary of DLD properties, including wave speed and domain size. **(g)** Apical vs. basolateral imaging reveals that both ATP and ACh sensors are activated specifically at the basolateral membrane following succinate (tuft cell) or high K⁺ (NE cell) stimulation, indicating basolateral secretion of both neurotransmitters. **(h)** Integrated model of airway environmental sensing and response through DLDs.

To definitively establish a relationship between ATP waves and DLD domain size, we developed an ATP/Ca^2+^ dual-sensor system (green ATP sensor, red XCaMPR). This revealed three distinct classes of succinate-induced DLDs (Fig. 4a, b, f and Supplementary Video 7). Type A DLDs, representing ∼50% of DLDs, were characterized by a small ATP wave with a surrounding larger calcium wave. Type B DLDs (∼33%) were characterized by a large calcium wave with no detectable ATP wave. Type C DLDs (∼18%) were characterized by a small calcium wave perfectly co-localizing with the ATP wave (Fig. 4a, b, Extended Data Fig. 11a). With *Chat KO*, only Type C DLDs were found (Fig. 4a). Thus, Types A and B DLDs require ACh diffusion, while Type C DLDs are purely ATP-dependent. Indeed, we demonstrated that tuft cells secrete ATP through the CALHM1 channel, as *Calhm1 KO*^38^ completely abolished Type A and Type C waves (Fig. 4a).

Given that NE cells have recently been shown to secrete ATP to activate myoepithelial cells in airway submucosal glands^10^ and vagal sensory nerves^9^, we hypothesized that ATP diffusion is responsible for NE cell-initiated DLDs. Indeed, high K^+^ triggered ATP waves (Extended Data Fig. 9b). Using the ATP/Ca^2+^ dual-sensor system, we found that high K^+^-induced waves of calcium activation were perfectly co-localized with ATP waves (Fig. 4d). Moreover, the P2 purinergic receptor antagonist suramin completely blocked these NE-initiated DLDs (Fig. 4e). Thus, both NE DLDs and Type A and C tuft DLDs rely on ATP diffusion. However, unlike tuft cells that secrete ATP through CALHM1, NE cells secrete ATP through a distinct mechanism, as *Calhm1 KO* does not affect NE cell-initiated DLDs (Extended Data Fig. 11b).

In summary, we have identified two neurotransmitters—ACh and ATP—whose diffusion governs the properties of DLDs, including their size and speed. Tuft cells can generate three distinct DLD patterns (A, B, C) based on the combination of ACh and ATP secretion, while NE cells generate a single, ATP-dependent DLD. DLDs that are dependent on ATP are both smaller and slower, while DLDs that rely on ACh are both larger and faster, independent of the initiating sentinel cell identity (Fig. 4f).

To determine whether ACh and ATP secretion occur at the apical or basolateral membrane, we employed high-speed, high-resolution spinning-disk confocal microscopy coupled with live sensors. Activation of ACh and ATP sensors occurs at the basolateral membrane (22 out of 22 for ATP; 39 out of 43 for ACh) (Fig. 4g, Extended Data Fig. 12, and Supplementary Video 8,9). We therefore hypothesize that neurotransmitters diffuse through the paracellular space, a continuous compartment underneath the tight junctions, thus shielded from the potentially disruptive luminal environment (Fig. 4h).

## Distinct DLD networks functionally respond to distinct stimuli, can be tuned for local or tissue-wide responses, and are patterned relative to functional organ anatomy

Having established the molecular and cellular basis of DLD activation, we next turned our attention to establishing how ensembles of sentinel cells are coordinated across the epithelium. We first sought to confirm that distinct tuft cell stimuli reproducibly activate the sentinel cell network in the same fashion. Surprisingly, the bitter compound denatonium, which activates taste receptor signaling in tuft cells^4–8,12^, resulted in a pattern of overlapping DLDs that allowed full coverage of the airway epithelium, but was distinct from the pattern of succinate-induced DLDs (Fig. 5a, left). We confirmed this finding in tracheal explants (Fig. 5a, right). This demonstrates that distinct distributed networks of tuft cells, with partially overlapping stimulus sensitivity, are positioned to ensure comprehensive epithelial coverage. In contrast, a local stimulus (1.5M KCl) visualized using a green fluorescent tracer, resulted in restricted local coverage (Fig. 5b and Supplementary Video 10). Titrating the stimulus concentration revealed that lower doses of either succinate or K⁺ resulted in attenuated DLDs, characterized by a reduction in number, domain size, and amplitude of calcium transients (Extended Data Fig. 13).

**Figure 5.**
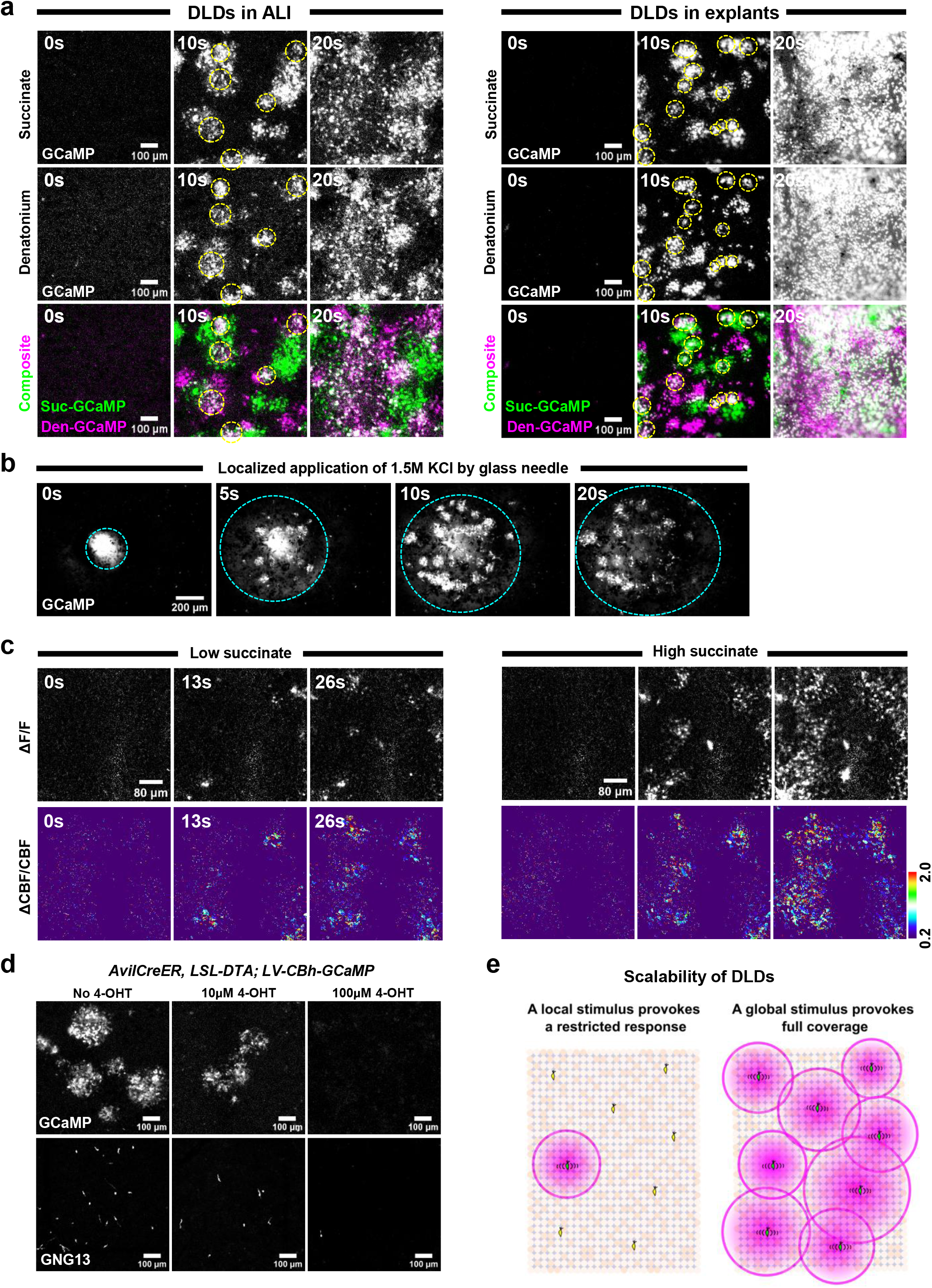
Distinct DLD networks functionally respond to distinct stimuli and can be tuned for local or tissue-wide responses. (**a**) Succinate and denatonium induce distinct patterns of overlapping DLDs that allow full coverage of the airway epithelium in both ALI cultures and explants. The 10s time point shows overlapping DLDs (yellow dashed circles) and DLDs exclusive to succinate (green) or denatonium (magenta). (**b**) Local administration of 1.5M KCl with a green fluorescent tracer evokes spatially restricted DLDs, confined to the area of tracer diffusion (cyan dashed circle) (**c**) Scaling of mucociliary output. Sequential application of low (200 µM) and high (2 mM) succinate to the same tracheal explants produces graded responses in both calcium activation (ΔF/F of GCaMP, top) and ciliary beat frequency (ΔCBF/CBF, bottom, pseudocolored). 200 µM succinate evokes sparse isolated DLDs with correspondingly limited tissue-wide CBF activation; 2 mM succinate evokes widespread DLDs with an accompanying broader amplification of CBF. (**d**) Partial genetic ablation of tuft cells using titrated 4-hydroxytamoxifen (4-OHT) in *AvilCreER; LSL-DTA; LV-CBh-GCaMP6s* ALI cultures progressively reduces DLD formation. Tuft cell density quantified by GNG13 staining. (**e**) Schematic model illustrating the scalable architecture of sentinel cell networks.

We next asked whether the relative activation of distinct sentinel cell networks scales to physiologic output. To do so, we sequentially applied varying concentrations of succinate to individual tracheal explants. Lower succinate concentration evoked sparse, localized CBF activation, whereas higher concentration evoked dense and widespread CBF activation (Fig. 5c). Furthermore, ablation of a limited number of tuft cells prevents the full coverage response (Fig. 5d, e). Thus, distinct NE and tuft cell sentinel cell networks translate graded sensory input into graded physiologic output, allowing either local or tissue-wide activation.

Furthermore, we find that sentinel cell networks can be spatially organized and vary with respect to anatomically localized functional organ compartments. Smooth muscle is localized to the posterior membrane of the trachea, while the remaining trachea is reinforced by non-contractile cartilage rings. This arrangement permits coughing without airway collapse. Both ACh and ATP are known to cause smooth muscle contraction. Thus, in principle, sentinel cell basolateral ACh and ATP secretion could evoke the contraction of smooth muscle. We confirmed that ACh induced robust smooth muscle contraction in tracheal explants. However, succinate application failed to do so (Extended Data Fig. 14 a,b and Supplementary Video 11). Examination of sentinel cell distribution along the airway revealed an anatomic basis for the lack of response. Both tuft cells and NE cells are diminished in the epithelium overlying smooth muscle (Extended Data Fig. 14 c,d). Thus, while DLDs determine zones of local physiologic activation within the epithelium itself, the anatomic distribution of DLD-inducing sentinel cells may pattern organ-level physiology.

## Disease-relevant cytokines enhance or limit DLDs and their corresponding physiologic output

Having established that DLDs are fundamental physiologic units, we sought to determine if and how they may be altered in disease states. Increased IL-1β levels have been shown in human patients with bronchiectasis^41^ and asthma^42^. Interestingly, IL-1β has been shown to enhance waves of calcium activation in astrocytes by increasing the number of cells responsive to purinergic signaling and upregulating P2Y receptor transcription^43^. Here, we found that treating ALI cultures with IL-1β significantly increased DLD intensity and size (Fig. 6a). To pinpoint the mechanism of this cytokine-induced alteration, we visualized neurotransmitter dynamics following cytokine application using live sensors. IL-1β potentiated the secretion of both ACh and ATP (Fig. 6d, f). Furthermore, using a *Chat-GFP* reporter^44^, we found that IL-1β upregulated the expression of *Chat* (Fig. 6e), suggesting that increased ACh synthesis underpins the DLD enhancement.

**Figure 6.**
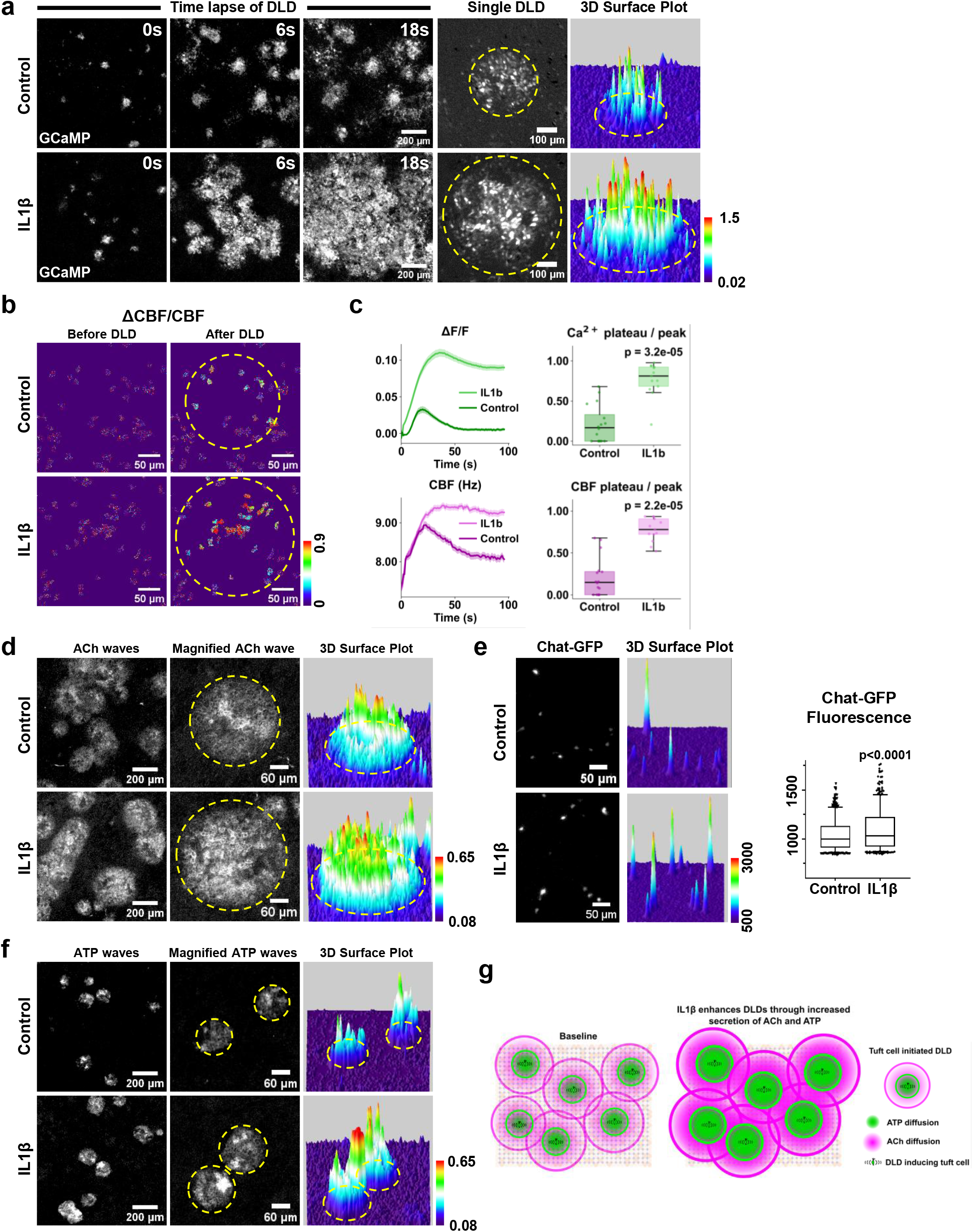
Disease-relevant cytokines enhance DLDs and their corresponding physiologic output. (**a**) IL-1β treatment enhances the intensity and domain size of succinate-induced DLDs in ALI cultures. Time-lapse imaging of calcium activation (GCaMP) shown for control (top) and IL-1β-treated (bottom) cultures. Right panels show a representative single DLD and a corresponding 3D surface plot of fluorescence intensity (yellow dashed circles outline the DLD). **(b)** Ciliary beat frequency changes (ΔCBF/CBF) imaged in the same field of view before and after succinate triggering in control and IL-1β-treated cultures. IL-1β-treated cultures show significantly expanded and intensified CBF activation. Yellow dashed circles denote the DLD domain. **(c)** Quantitative traces of calcium (ΔF/F, top) and CBF (Hz, bottom) over time in control vs. IL-1β-treated cultures. Shaded bands denote SEM. IL-1β increases both the peak amplitude and the sustained plateau (average over 66-96s time window) of both signals. Box plots show the plateau-to-peak ratio for calcium and CBF, with significant elevation in the IL-1β condition (Mann-Whitney U test). Boxes show median and interquartile range; whiskers extend to 1.5×IQR. **(d)** IL-1β potentiates succinate-induced ACh secretion. ACh sensor signal shown for control (top) and IL-1β-treated (bottom) cultures and corresponding 3D surface plots (yellow dashed circles outline DLDs). **(e)** IL-1β upregulates *Chat* expression in tuft cells, measured by *Chat-GFP* reporter fluorescence. Representative images shown with corresponding 3D surface plots and quantification of Chat-GFP⁺ cells (Mann-Whitney U test; whiskers represent 5th-95th percentile). **(f)** IL-1β potentiates succinate-induced ATP secretion. ATP sensor signal shown for control (top) and IL-1β-treated (bottom) cultures and corresponding 3D surface plots (yellow dashed circles outline DLDs). **(g)** Schematic model of IL-1β-mediated DLD remodeling, illustrating amplification of DLD size and intensity.

We next assessed the physiologic relevance of IL-1β-induced DLD enhancement. IL-1β treatment resulted in an amplification of succinate-induced CBF responses (Fig. 6b). Both peak amplitude and the plateau-to-peak ratio were elevated for both calcium and CBF (Fig. 6c). Therefore, IL-1β addition resulted in a sustained synchronized increase of calcium and CBF.

We next bypassed natural ligand sensing through optogenetic tuft cell activation. IL-1β-induced enhancement of DLD calcium and CBF responses were preserved (Extended Data Fig. 15), demonstrating that IL-1β action is independent of the succinate stimulus.

In direct contrast to IL-1β, treatment with IFN-γ, critical for antiviral defense, significantly reduced the density of succinate-induced DLDs (Extended Data Fig. 16a). We hypothesized that this could result from a loss of sentinel cells themselves. Indeed, quantification of tuft cells revealed a significant decrease in their numbers following IFN-γ treatment (Extended Data Fig. 16). In summary, pro-inflammatory cytokines can remodel DLDs and their distribution both positively and negatively with corresponding physiologic consequences (Fig. 6g, Extended Data Fig. 16b).

## Pathogen sensing evokes DLDs

Having defined the architecture of DLDs and their modulation by disease-relevant cytokines, we sought to determine whether DLDs sense complex environmental perturbations of clinical relevance. The mechanisms for sensing pathogenic fungi are understudied, despite the fact that invasive fungi are a leading cause of invasive pulmonary infection in immunocompromised patients^45^. We previously demonstrated that *Aspergillus*-derived melanin, a major constituent of the fungal cell wall, suppresses airway epithelial calcium signaling and chemokine secretion, revealing that the fungus actively subverts epithelial defenses^46^. Here we asked whether the airway epithelium possesses a reciprocal sensing mechanism capable of detecting the presence of the fungus. Therefore, we measured airway epithelial calcium responses to conditioned media (CM) from *Aspergillus fumigatus* cultures. *Aspergillus* CM robustly induced DLDs in both air-liquid interface (ALI) cultures and tracheal explants (Fig. 7a). We next established that these DLDs emanated from tuft cells (Fig. 7b), which are known to sense bacteria and protists. Indeed, genetic ablation of tuft cells abrogated DLDs (Fig. 7c). To characterize the molecular basis of sensing, we performed heat inactivation and molecular size fractionation (Fig. 7d). Heat inactivation did not alter DLD formation ruling out a heat-labile protein. We next fractionated the CM using a 3-kDa molecular weight cutoff filter. The protein-depleted small-molecule-containing fraction (<3 kDa + small molecules) fully preserved DLD-inducing activity; the high molecular weight protein fraction (>3 kDa) failed to induce DLDs.

**Figure 7.**
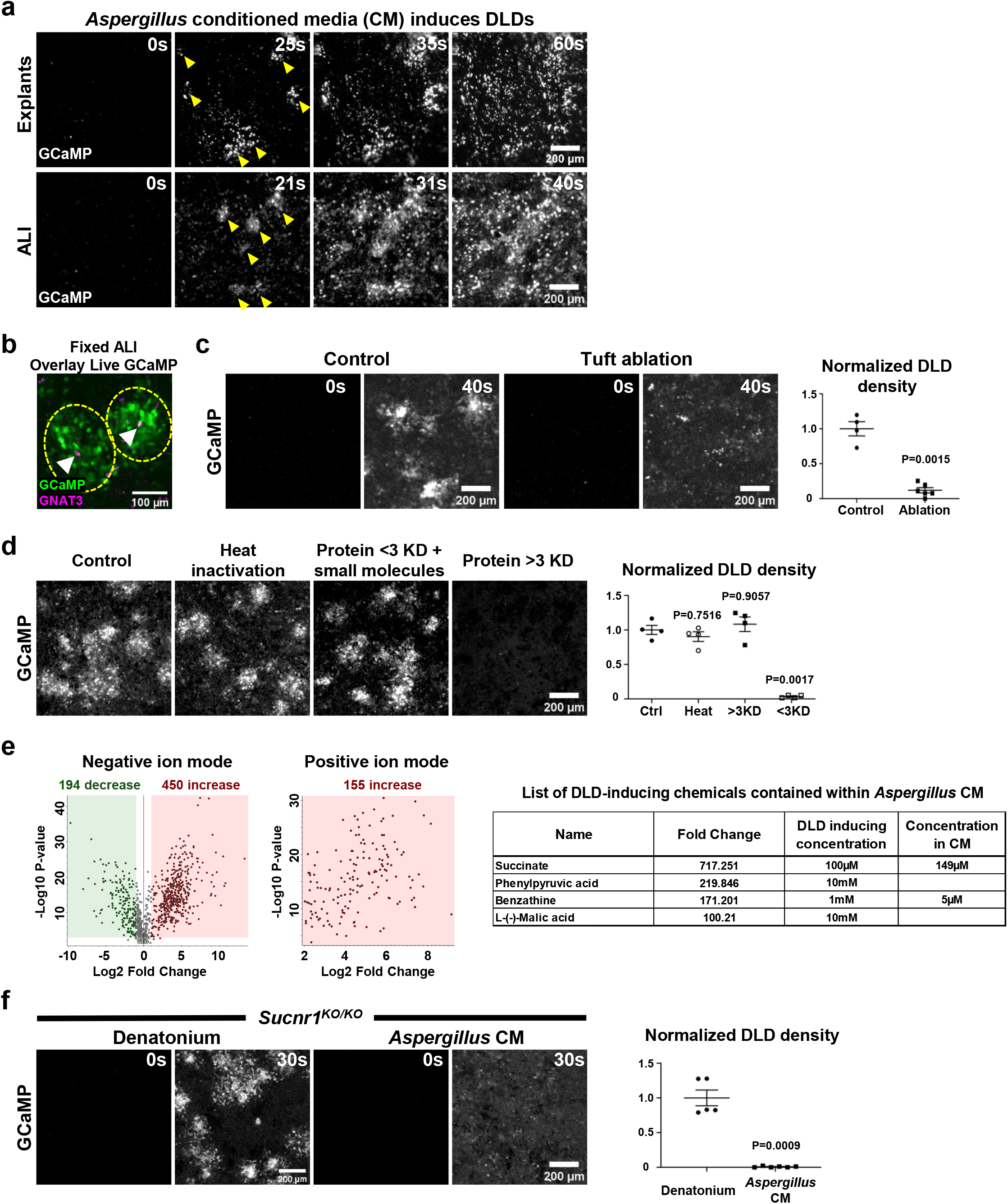
SUCNR1-mediated detection of a fungal metabolite is required for *Aspergillus fumigatus*-induced DLDs. **(a)** *Aspergillus fumigatus* conditioned media (CM) induces DLDs in both tracheal explants (top) and ALI cultures (bottom). Yellow arrowheads indicate emerging foci of activation at early time points. **(b)** Post-imaging immunostaining identifies GNAT3⁺ tuft cells (white arrowheads) at the centers of *Aspergillus* CM-induced DLDs. Yellow dashed circles denote DLD domains. **(c)** Genetic ablation of tuft cells (*AvilCreER; LSL-DTA*) abrogates *Aspergillus* CM-induced DLDs in ALI cultures. Time-lapse calcium imaging shown for control and tuft cell-ablated cultures (left); normalized DLD density quantified at right (Welch’s t-test). **(d)** Characterization of the active component of *Aspergillus* CM. Heat inactivation fails to reduce DLD-inducing activity. Size fractionation employing a 3 kDa molecular weight cutoff filter shows that activity is preserved in the small-molecule fraction (<3 kDa) and absent in the protein fraction (>3 kDa). Normalized DLD density quantified at right (one-way ANOVA with Games-Howell post-hoc test). **(e)** Untargeted LC-HRMS profiling of *Aspergillus* CM relative to uninoculated control media. Volcano plots show metabolites detected in negative ion mode (left) and positive ion mode (middle) (significance thresholds: p < 0.001, log₂ fold change > 1). The table at right lists candidate DLD-inducing metabolites identified from the most enriched compounds. Of the four active candidates, only succinate induces DLDs at concentrations within its physiologic range in CM. **(f)** *Aspergillus* CM does not induce DLDs in *Sucnr1*⁻/⁻ epithelium. Denatonium, a bitter taste receptor agonist that activates tuft cells through an independent pathway, continues to induce DLDs. Normalized DLD density quantified at right (Welch’s t-test).

To identify DLD-inducing small molecules, we employed untargeted liquid chromatography–high resolution mass spectrometry (LC-HRMS) in both negative and positive electrospray ionization modes on *Aspergillus* CM to comprehensively profile the fungal exometabolome (Fig. 7e). In negative ionization mode, a clear bidirectional metabolic shift was observed, characterized by 450 increased and 194 decreased chemical features (Fig. 7e). Conversely, positive ionization mode captured exclusively upregulated features (155 increased, 0 decreased; Fig. 7e). We next screened metabolites enriched more than 100-fold relative to control media to test their ability to induce DLDs (Extended Data Fig. 17). Among the 18 tested highly enriched candidates, only 4 produced DLDs (Fig. 7e, Extended Data Fig. 17). Of these, only succinate was able to induce DLDs at a physiologically relevant micromolar concentration. We therefore hypothesized that *Aspergillus*-derived succinate is sensed by tuft cells via the succinate receptor SUCNR1, whose expression is uniquely enriched in tuft cells and is known to participate in sensing bacteria and protists. Remarkably, *Aspergillus* CM failed to induce any DLDs in Sucnr1 KO epithelium (Fig. 7f). Together, these results demonstrate that airway tuft cells detect *Aspergillus*-derived succinate through SUCNR1 to induce DLDs.

## Discussion

The airway epithelium is the first line of defense against inhaled environmental stimuli and pathogens. Inhaled stimuli are known to activate sentinel cells which communicate with sub-epithelial immune cells and vagal sensory afferents using neurotransmitters such as ACh and ATP^3–12^. Herein, we define a very simple model in which diffusion of these same sentinel cell-derived neurotransmitters results in activation of circular diffusion-limited domains (DLDs) of neighboring secretory, ciliated, and stem cells. In our model, neurotransmitter secretion and diffusion dictate the speed, intensity, and domain size of DLDs. The basolateral locus of secretion allows for the diffusion of neurotransmitters through both the local paracellular epithelial space and across the basement membrane to reach immune cells and vagal sensory afferents^3–12^. Thus, neurotransmitter diffusion represents the basis for the activation of both local epithelial responses and organismal responses mediated by neural circuitry^9^.

We show that DLDs represent a general mechanism for patterning physiologic domains of the airway epithelium irrespective of the identity of the underlying sentinel cells. In fact, we establish that the airway possesses a diverse functional repertoire of sentinel cells, including tuft cells that secrete ACh, ATP, or both, and NE cells that secrete ATP through a CALHM1-independent mechanism distinct from the tuft cell CALHM1 channel. Furthermore, the distribution of these diverse sentinel cells leads to the formation of distinct sensory networks that are responsive to distinct stimuli and insulated from one another, as tuft cell-initiated DLDs do not result in the activation of NE DLDs, nor do NE cell-initiated DLDs provoke tuft cell DLDs. Additionally, sentinel cells are distributed in a manner that is sufficient to ensure comprehensive epithelial coverage. However, the amplitude and number of DLDs can be scaled according to stimulus intensity, which in turn permits locally restricted responses. Furthermore, the distribution of sentinel cells varies with tissue anatomy, limiting their effects on target cells. For example, the lack of sentinel cells in the posterior membranous tracheal epithelium precludes robust smooth muscle activation. However, since sentinel cell populations are known to be expanded in diseases of the airway, alteration in their distribution may engender aberrant smooth muscle activation.

We show that DLDs are associated with both cardinal epithelial physiologic functions: mucociliary transport and epithelial renewal through stem cell proliferation. Although airway stem cells were not previously known to be participants in sentinel cell-induced calcium waves, spontaneous calcium waves have been associated with radial glial cell proliferation during neocortex development^47^ and with cell-cycle progression in adult epidermal basal stem cells^48^. Our findings reveal a complementary mechanism in which stem cell replication is driven by activation of sentinel cells. Furthermore, inflammatory cytokines are shown to remodel DLDs. IL-1β augments DLDs by potentiating neurotransmitter secretion, while IFN-γ dampens DLDs by depleting the sentinel cell pool itself, demonstrating that cytokines can either augment or dampen protective physiologic responses.

Finally, we show that pathogenic fungi provoke DLDs. Interestingly, succinate sensing by tuft cells has also been documented to induce protective type 2 inflammation in protist infections^49,50^. Thus, both local physiologic and broader inflammatory protective responses to pathogens are elicited by succinate-triggered tuft cells. Furthermore, our findings take on additional significance in light of our prior demonstration that *Aspergillus* melanin potently suppresses airway epithelial calcium fluxing and downstream chemokine secretion, thereby reducing neutrophil recruitment to the airway^46^. Thus, it would appear that tuft cell calcium signaling represents a critical node in governing host-pathogen interactions at the airway interface. The existence of both a fungal sensing mechanism and a fungal evasion strategy centered on the same epithelial calcium signaling pathways suggests that modulation of this molecular apparatus may be of therapeutic value.

In summary, we illustrate a set of design principles employed by sophisticated tissue-based local sensory networks. The sentinel cells themselves link differing environmental stimuli to local airway physiologic responses that can be scaled according to stimulus identity, modulated by disease-relevant cytokines, and sense real-world threats. Thus, we describe a unified fundamental model for airway sensing and response.

## Supporting information

Supplementary Video 1. Succinate induces Intercellular calcium waves (ICWs) in ALI culture

Supplementary Video 2. Chemogenetic activation of tuft cells induces robust ICWs

Supplementary Video 3. Optogenetic activation of tuft cells induces robust ICWs

Supplementary Video 4. High K+ induces ICWs in ALI culture

Supplementary Video 5. ACh diffusion delimits tuft cell-initiated ICW domain size

Supplementary Video 6. ICWs propagate across physical cuts

Supplementary Video 7. Tuft cells generate three types of DLDs

Supplementary Video 8. Tuft cells secrete ACh from the basolateral membrane

Supplementary Video 9. Tuft cells secrete ATP from the basolateral membrane

Supplementary Video 10. Local 1.5M KCl administration evokes spatially restricted DLDs

Supplementary Video 11.Succinate fails to induce robust smooth muscle contraction

---

**Extended Data Fig. 1.**
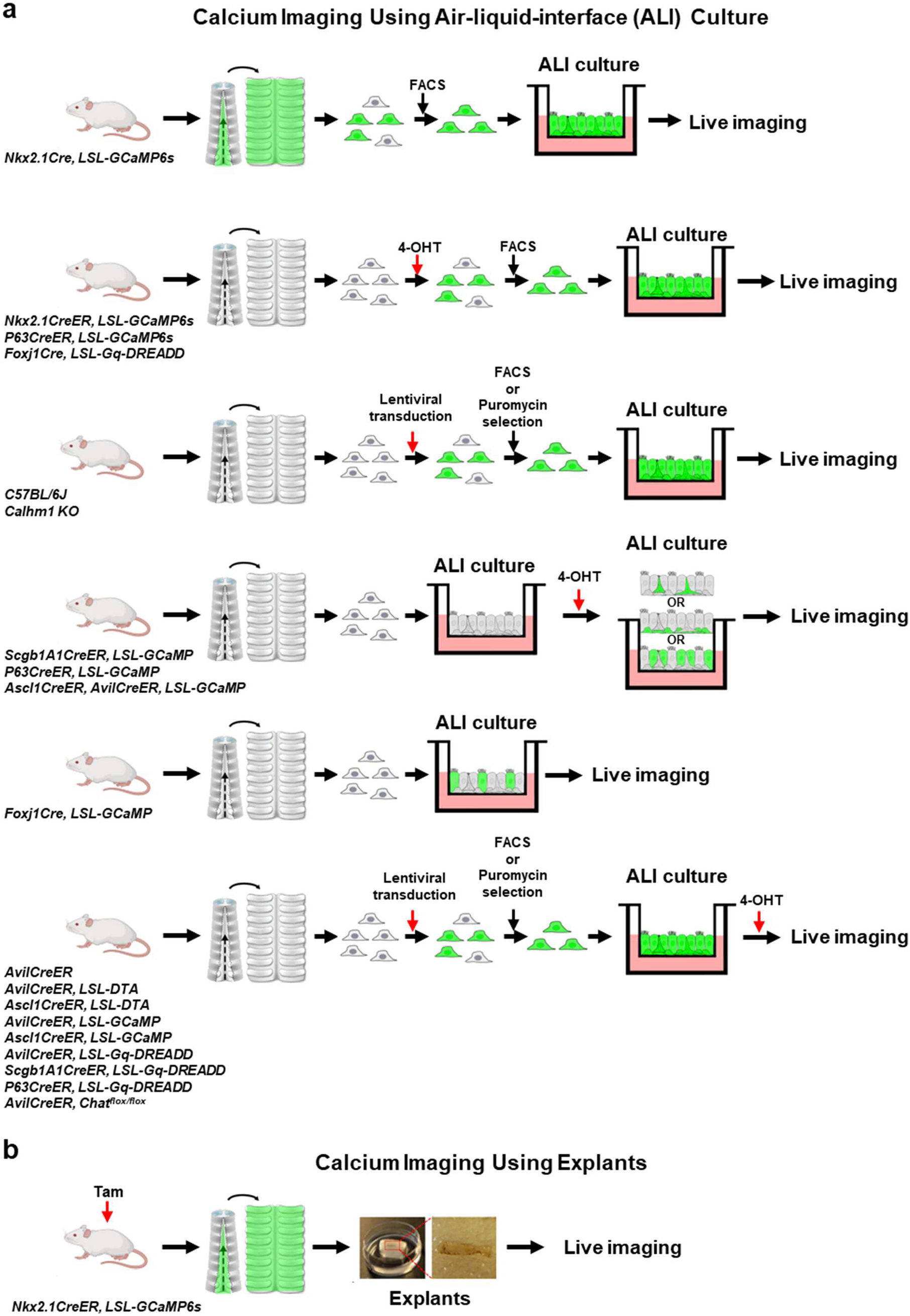
Experimental schemes for live calcium imaging. Live calcium imaging is performed with air-liquid-interface (ALI) cultures (**a**) or tracheal explants (**b**). Diagrams illustrate the genotypes of the mouse lines and the procedural workflow.

**Extended Data Fig. 2.**
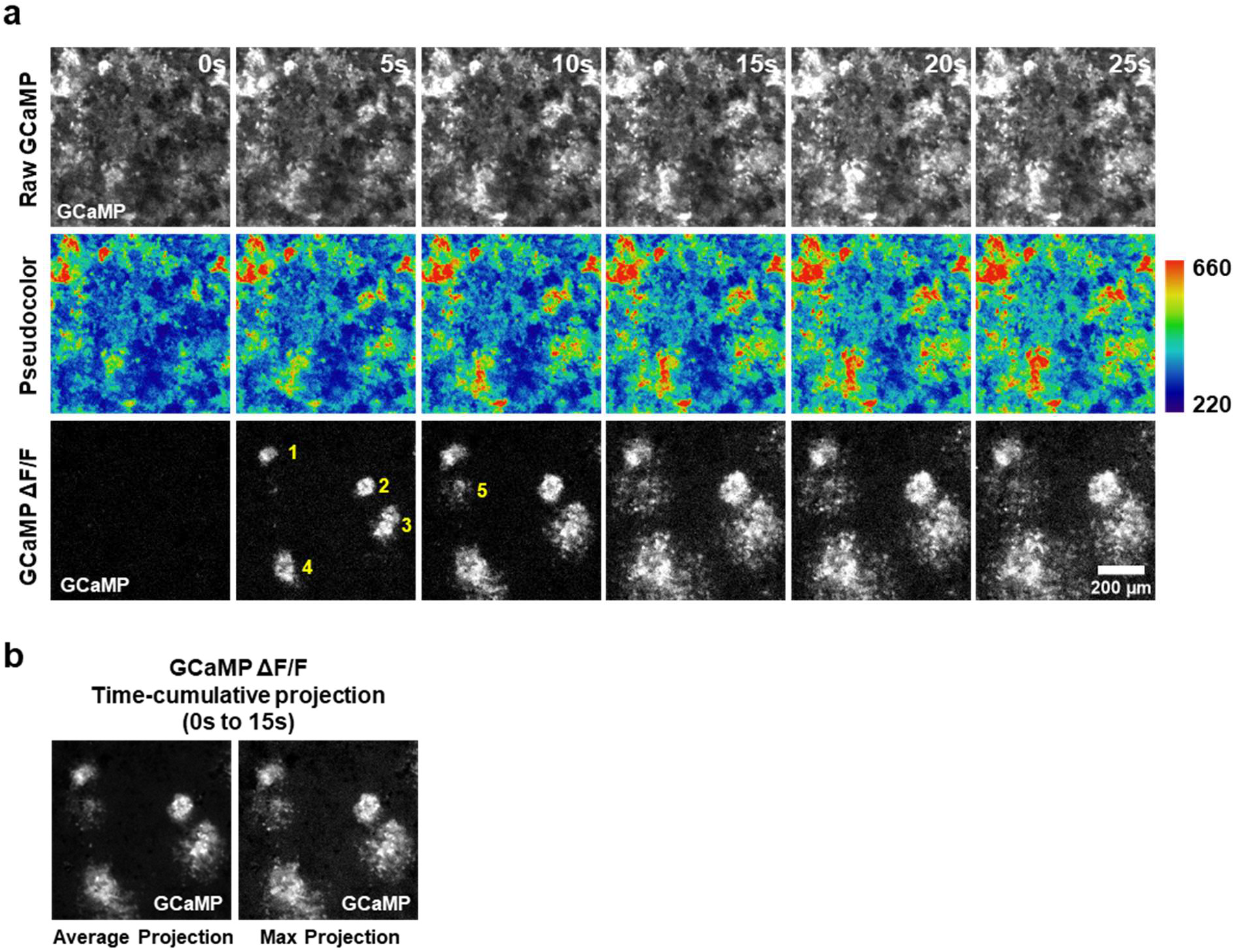
Presentation of live calcium imaging results. (**a**) Comparison of three methods for displaying succinate-induced ICWs in ALI cultures. From top to bottom: raw GCaMP fluorescence, pseudocolored GCaMP fluorescence, and ΔF/F (change in fluorescence over baseline). Yellow numbers in the bottom panel mark the emergence of individual ICWs. The ΔF/F method most clearly visualizes wave initiation and propagation and is used for all multi-timepoint representations of ICWs in this paper. (**b**) Time-cumulative projection of the ΔF/F data from (a). ΔF/F GCaMP fluorescence data acquired over a period of 15 seconds is projected into a single image using either average or maximum intensity projection. Both methods effectively capture the induction of ICWs. Time-cumulative projections are used for all single-image representations of ICWs in this paper.

**Extended Data Fig. 3.**
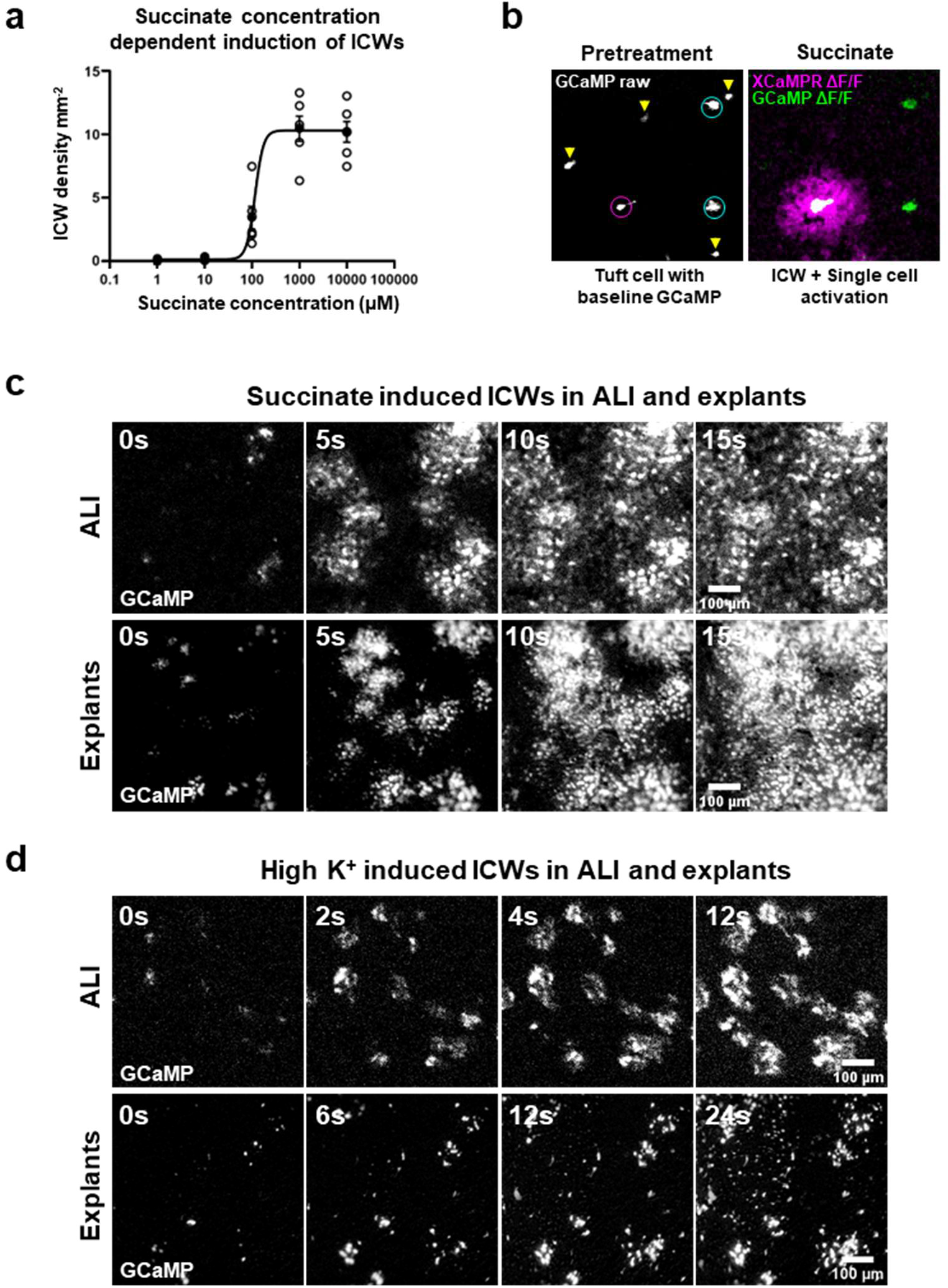
Characterization of tuft cell and NE cell-initiated ICWs. (**a**) Concentration-response curve for succinate-induced ICWs (EC₅₀ = 116 µM). (**b**) Functional heterogeneity of tuft cells in response to succinate. The magenta circle highlights a succinate-activated tuft cell (green) that initiates an ICW (magenta). Cyan circles indicate activated tuft cells that do not initiate ICWs. Yellow arrowheads mark tuft cells that are unresponsive to succinate. (**c**) Succinate induces ICWs in ALI and explants that originate from discrete foci and eventually cover the entire tissue. (**d**) High K^+^ induces ICWs in ALI and explants, with less expansive coverage.

**Extended Data Fig. 4.**
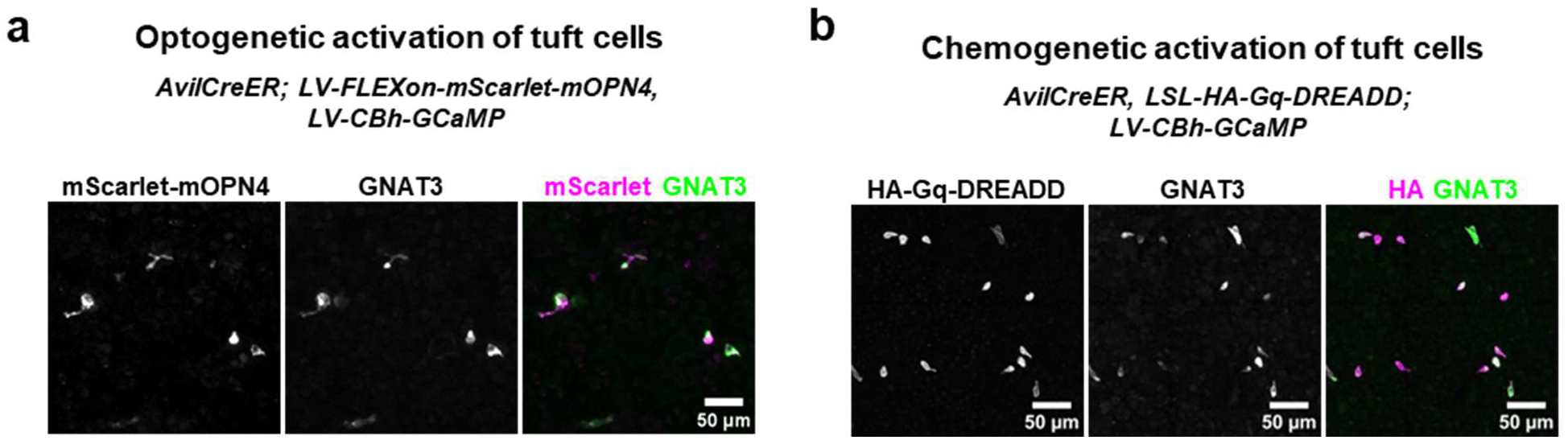
Validation of tuft cell-specific expression of optogenetic and chemogenetic actuators. (**a**) Co-localization of mScarlet-tagged optogenetic actuator mOPN4 and tuft cell marker GNAT3 confirms tuft cell specific expression. (**b**) Co-localization of HA-tagged chemogenetic actuator Gq-DREADD and GNAT3 confirms tuft cell-specific expression.

**Extended Data Fig. 5.**
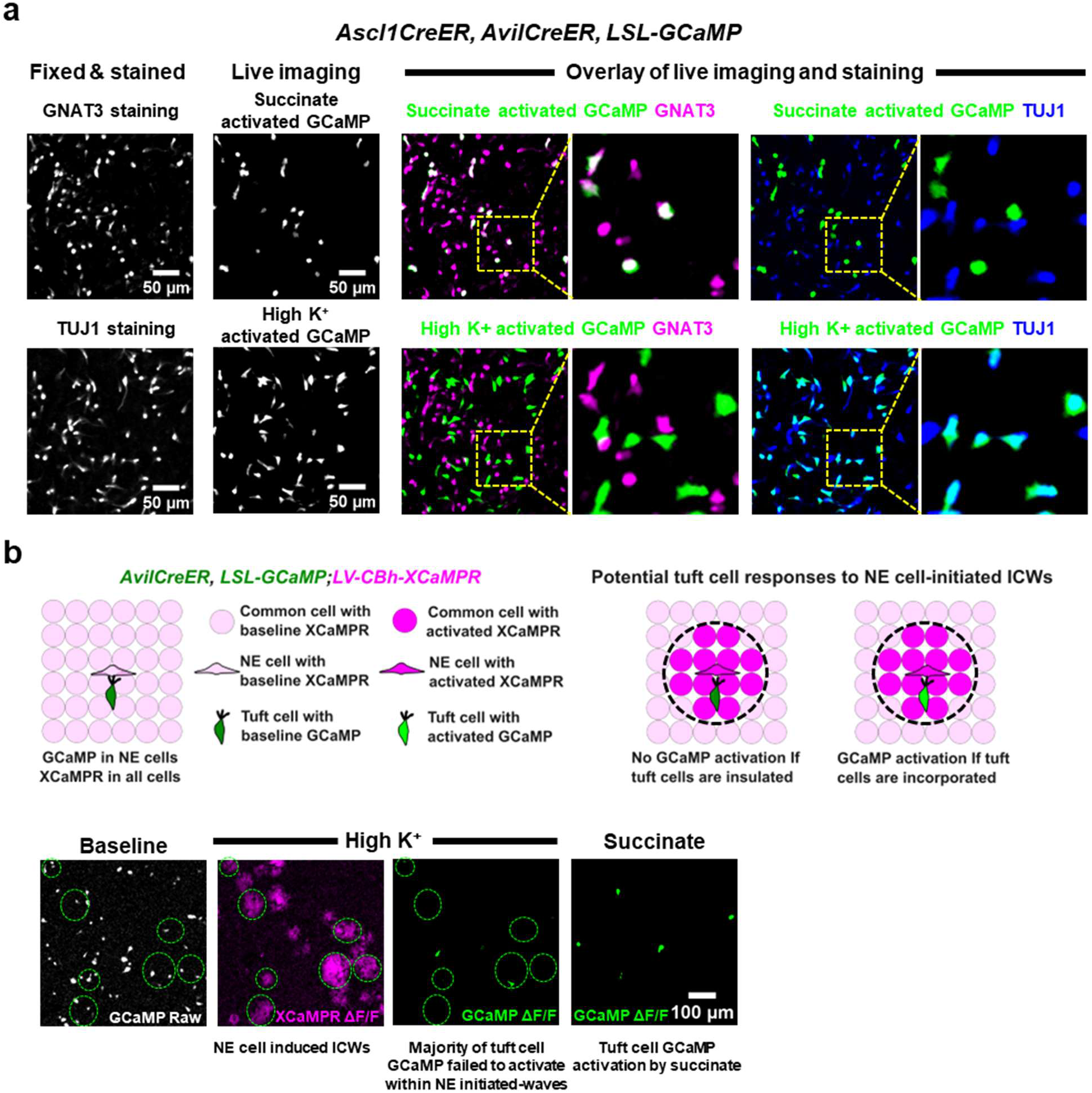
Tuft cells and NE cells are functionally insulated from each other. (**a**) Overlay of live calcium imaging with fixed immunostaining shows that succinate selectively activates GNAT3⁺ tuft cells, while high K⁺ selectively activates TUJ1⁺ NE cells. (**b**) The majority of tuft cells located within domains of NE cell-initiated ICWs (demarcated by green dashed circles in live imaging and black circles in schematics) do not exhibit cacium spikes, indicating that tuft cells are insulated from NE cell-initiated ICWs.

**Extended Data Fig. 6.**
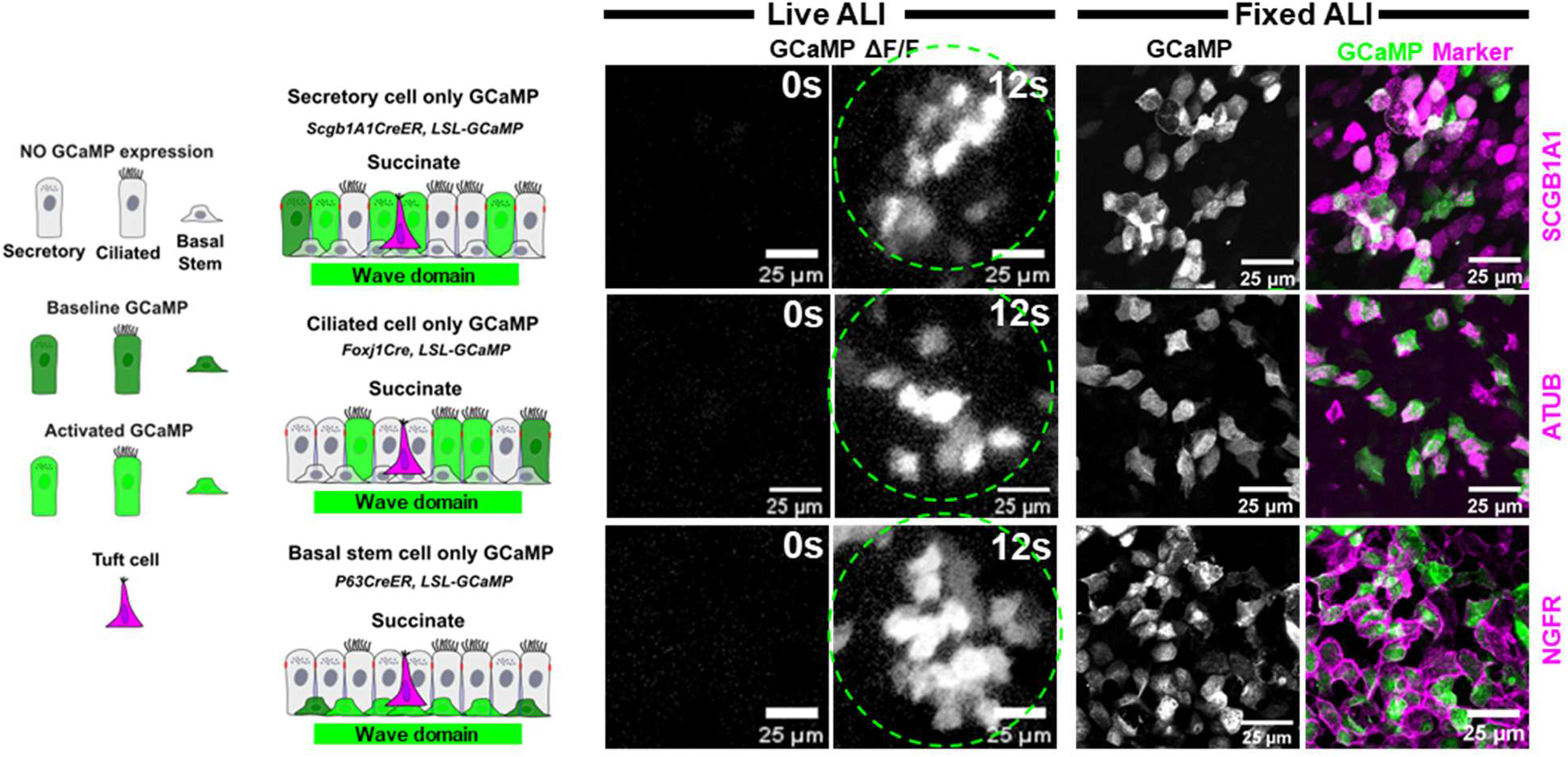
All common cells participate in tuft cell-initiated ICWs. Calcium imaging with cell type specific GCaMP expression reveals that succinate-induced ICWs activate secretory, ciliated, and basal stem cells, indicating that all three common cell types participate in tuft cell-initiated ICWs. Cell identities are confirmed by post-hoc immunostaining for common cell markers. The green dashed circles denote the wavefront of the propagating ICW. Time stamp in seconds shown at top right corner.

**Extended Data Fig. 7.**
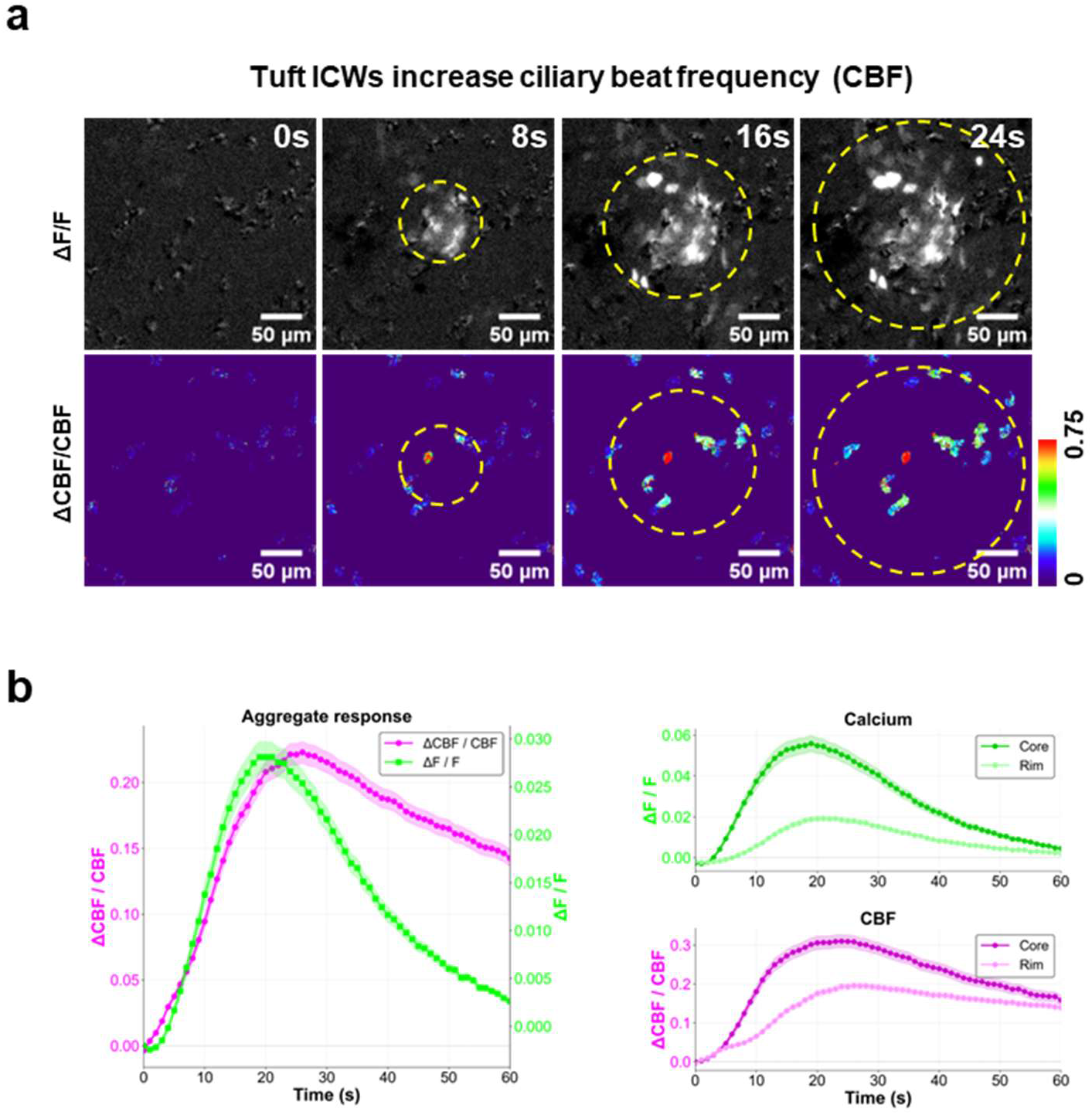
Stimulus-triggered tuft cell generate local domains of increased ciliary beat frequency. **(a)** High-speed live imaging of succinate-induced, tuft cell-initiated ICWs. Top row: calcium activation visualized as ΔF/F of the GCaMP signal within the ICW domain (yellow dashed circle). Bottom row: ciliary beat frequency change (ΔCBF/CBF) computed from the WGA-488 cilia signal, showing spatial coupling of ciliary activation to the calcium wave. Time stamps in seconds shown at top right. **(b)** Quantitative traces of calcium (ΔF/F, green) and CBF (ΔCBF/CBF, magenta) within tuft cell-initiated ICWs. Left: aggregate response across the entire ICW domain. Right: separation of responses into core (cells within half the wave radius) and rim (cells beyond half the wave radius) compartments, showing larger amplitudes in the core for both calcium and CBF. Together with the NE cell data shown in Fig. 2e, f, these results demonstrate that CBF coupling is a general property of sentinel cell-initiated functional domains irrespective of the initiating cell type.

**Extended Data Fig. 8.**
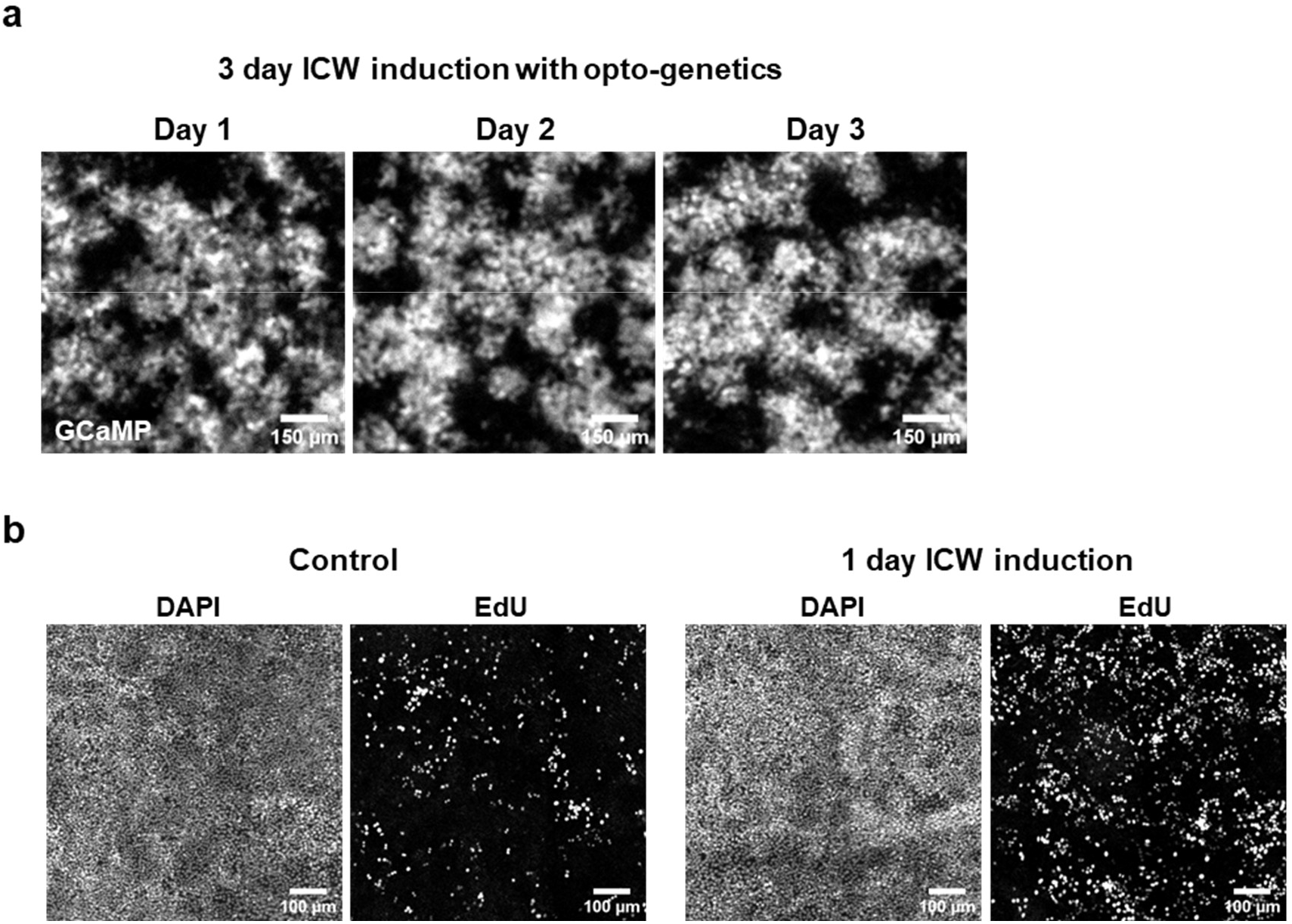
Optogenetically activated tuft cells drive stem cell proliferation. **(a)** Three day optogenetic ICW induction in *AvilCreER; LV-FLEXon-mOPN4; LV-CBh-GCaMP8s* ALI cultures. Representative GCaMP calcium imaging captured on each day of the induction protocol (Day 1, Day 2, Day 3) shows that robust ICWs are consistently induced over the three days of the chronic activation paradigm used in Fig. 2g, h. **(b)** One day optogenetic induction is sufficient to drive basal stem cell proliferation. DAPI and EdU staining in control (no mOPN4, left) and optogenetically activated (right) cultures. EdU⁺ cell density quantification shown in Fig. 2h.

**Extended Data Fig. 9.**
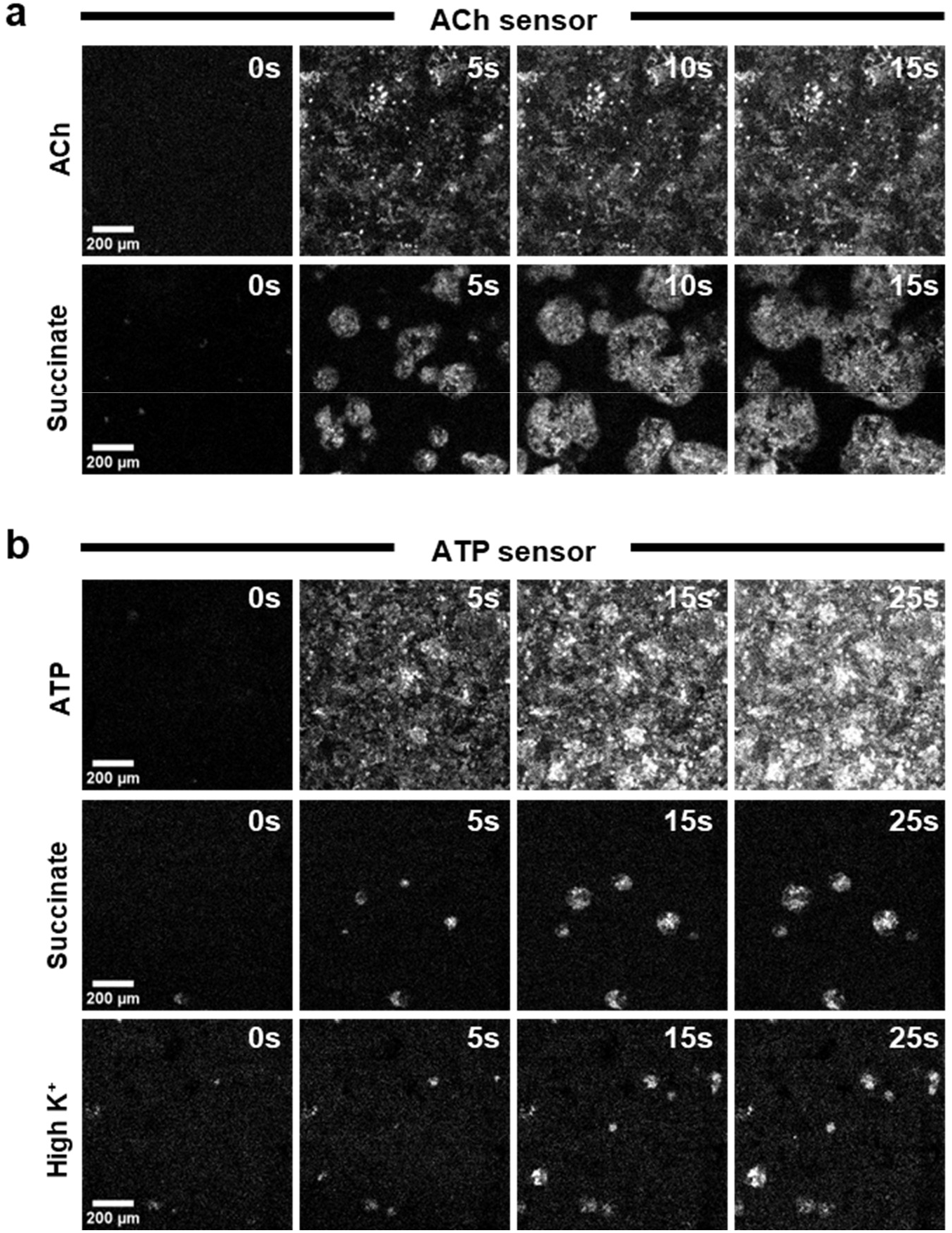
Sentinel cell stimuli activate ACh and ATP sensors. (**a**) ACh activates ACh sensors all at once whereas succinate induces ACh waves that originate from distinct foci and spread radially, similar to calcium waves. (**b**) ATP activates ATP sensors all at once whereas succinate and high K^+^ induce ATP waves that originate from distinct foci and spread radially, similar to calcium waves. Time stamp in seconds shown at top right corner.

**Extended Data Fig. 10.**
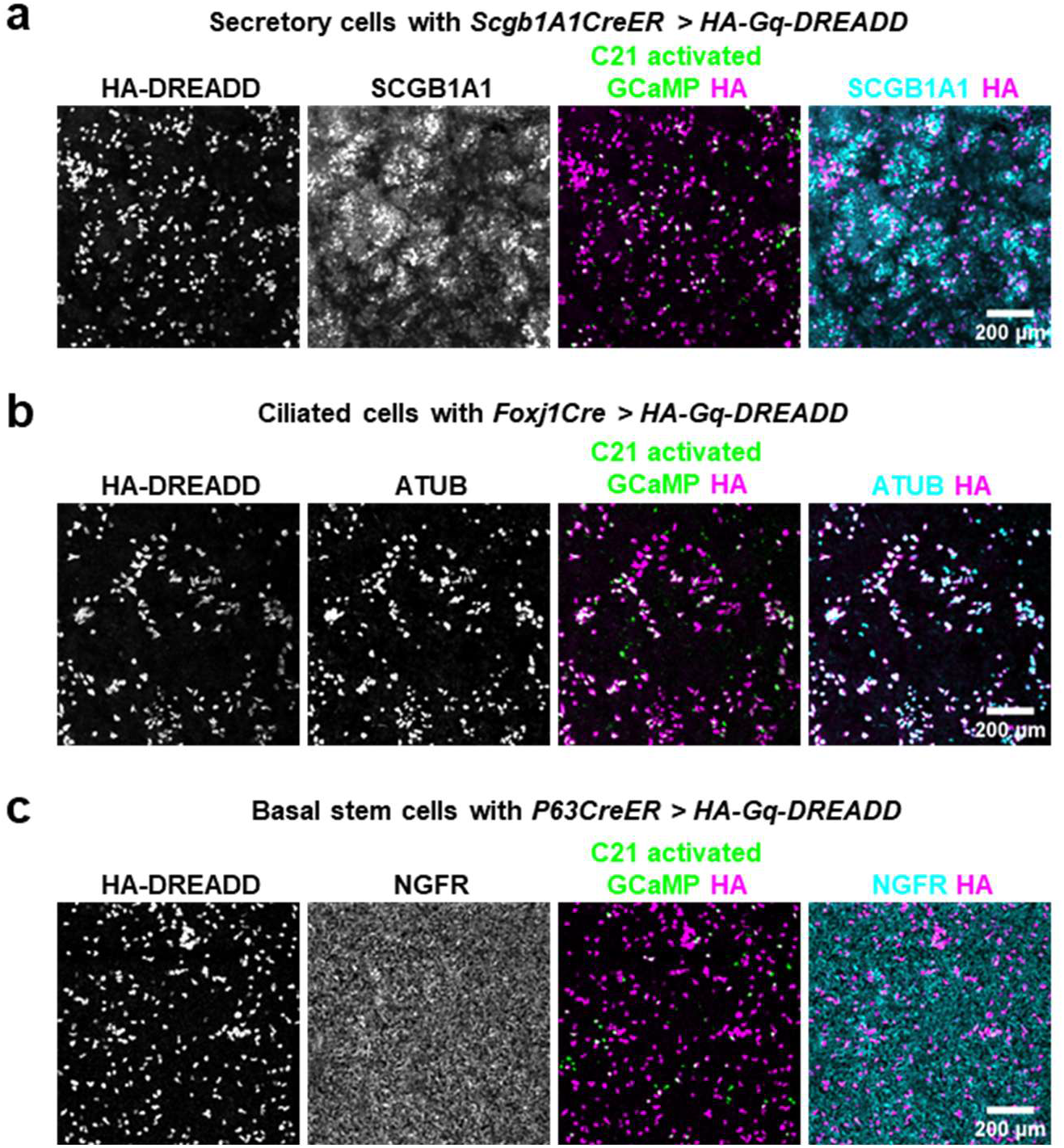
Validation of sparse chemogenetic activation within each common cell type. (**a**) Low dose (20nM) 4-Hydroxytamoxifen (4-OHT) induces sparse expression of HA tagged Gq-DREADD in SCGB1A1+ secretory cells using *Scgb1A1CreER* driver. (**b**) *Foxj1Cre* drives expression of HA tagged Gq-DREADD in all ATUB+ ciliated cells. (**c**) Low dose (10nM) 4-OHT induces sparse expression of HA tagged Gq-DREADD in NGFR+ basal stem cells using *P63CreER* driver. In all panels, DREADD ligand C21 induces calcium spikes (GCaMP, green) specifically in a subset of DREADD-expressing cells (HA, magenta).

**Extended Data Fig. 11.**
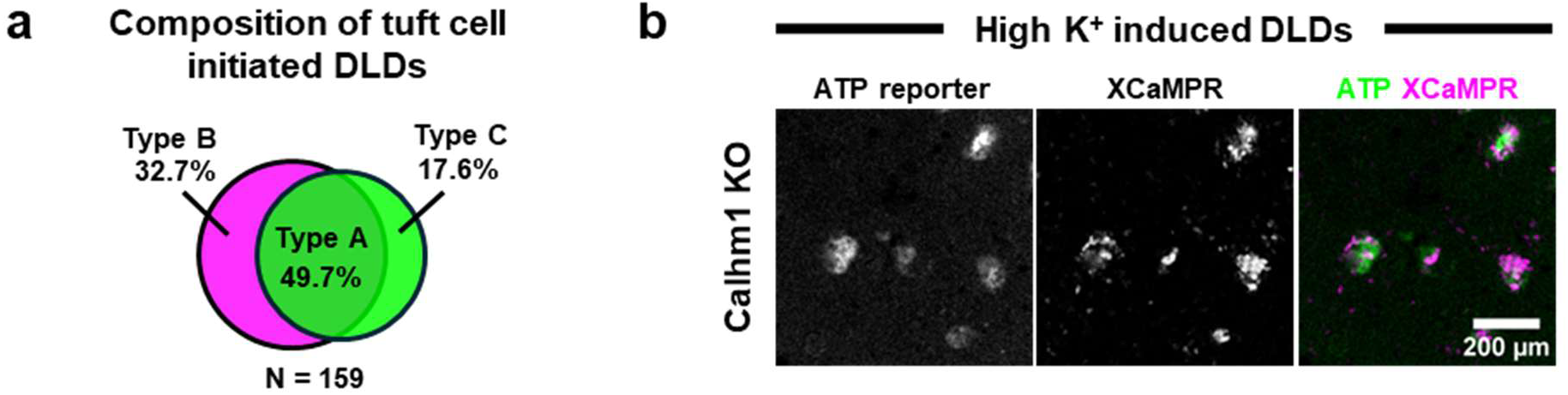
Characterization of DLD diversity. (**a**) Composition of three types of tuft cell-initiated DLDs induced by succinate. Green signifies use of ATP diffusion and magenta signifies use of ACh diffusion. (**b**) High K^+^ induced DLDs are independent of the ATP secretion channel CALHM1 as their formation is unaffected in *Calhm1* KO ALI cultures.

**Extended Data Fig. 12.**
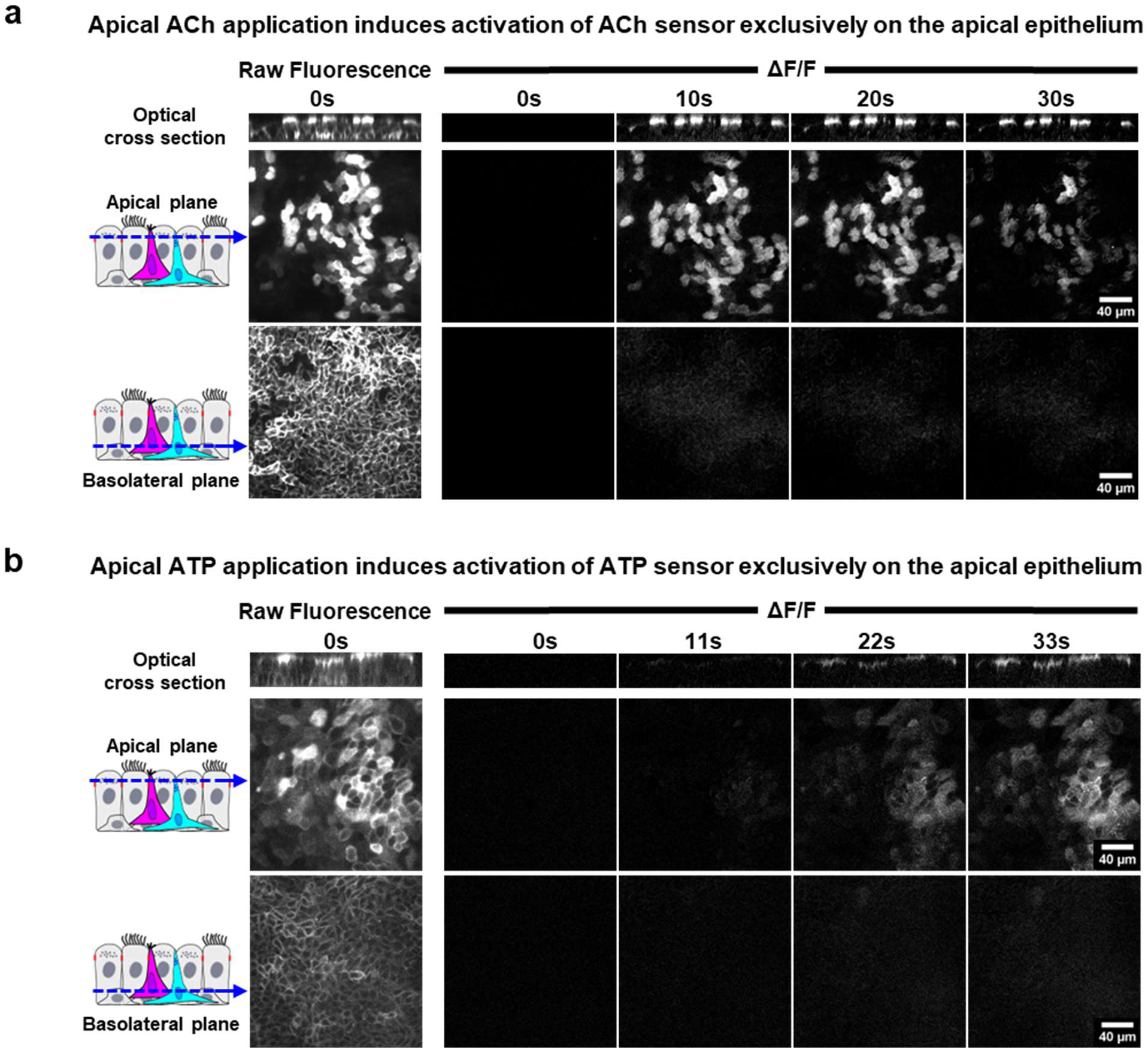
Apical ACh and ATP application results in apical ACh and ATP sensor activation without basolateral activation. (**a**) Apical ACh application induces activation of ACh sensor exclusively at apical side of epithelium. (**b**) Apical ATP application induces activation of ATP sensor exclusively at apical side of epithelium.

**Extended Data Fig. 13.**
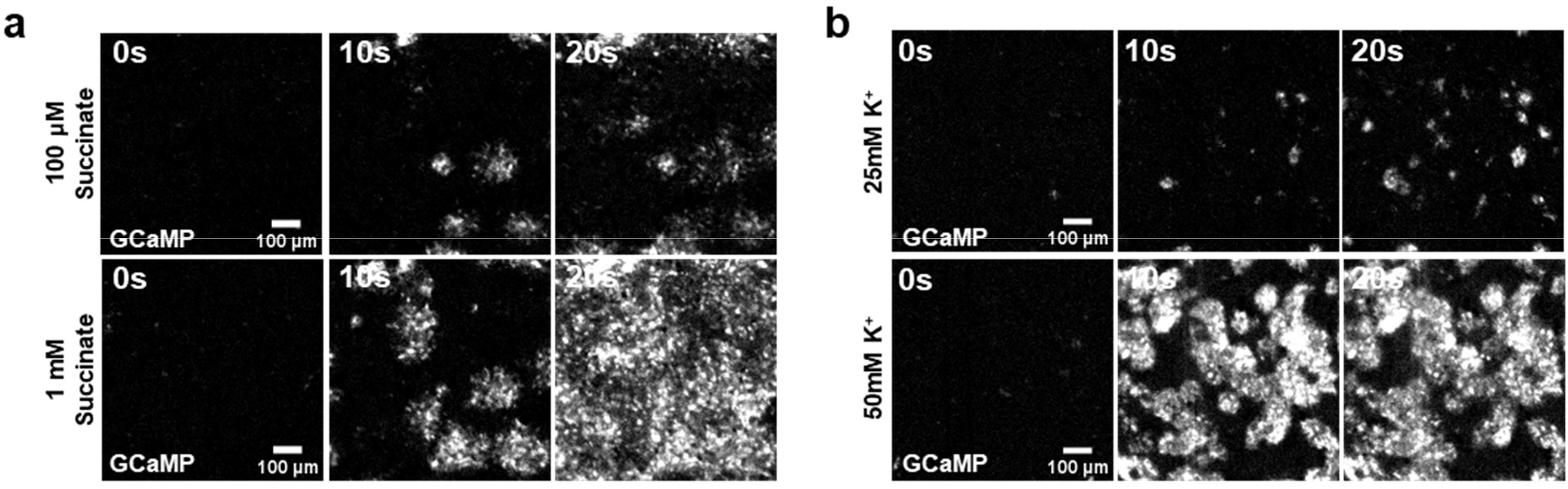
Stimulus strength scales with epithelial DLD intensity and coverage. (**a**) Time-lapse calcium imaging (GCaMP) of ALI cultures stimulated with low (100 µM) or high (1 mM) succinate. Low succinate evokes sparse DLDs with limited calcium activation; high succinate evokes dense, widespread DLDs across the same culture. (**b**) Time-lapse calcium imaging (GCaMP) of ALI cultures stimulated with low (25 mM) or high (50 mM) K⁺. Low K⁺ evokes fewer NE cell-initiated DLDs with smaller domains; high K⁺ evokes denser and larger DLDs. Together, these results demonstrate that DLD number, size, and calcium amplitude scale with stimulus intensity in both tuft cell-initiated (succinate) and NE cell-initiated (high K⁺) responses.

**Extended Data Fig. 14.**
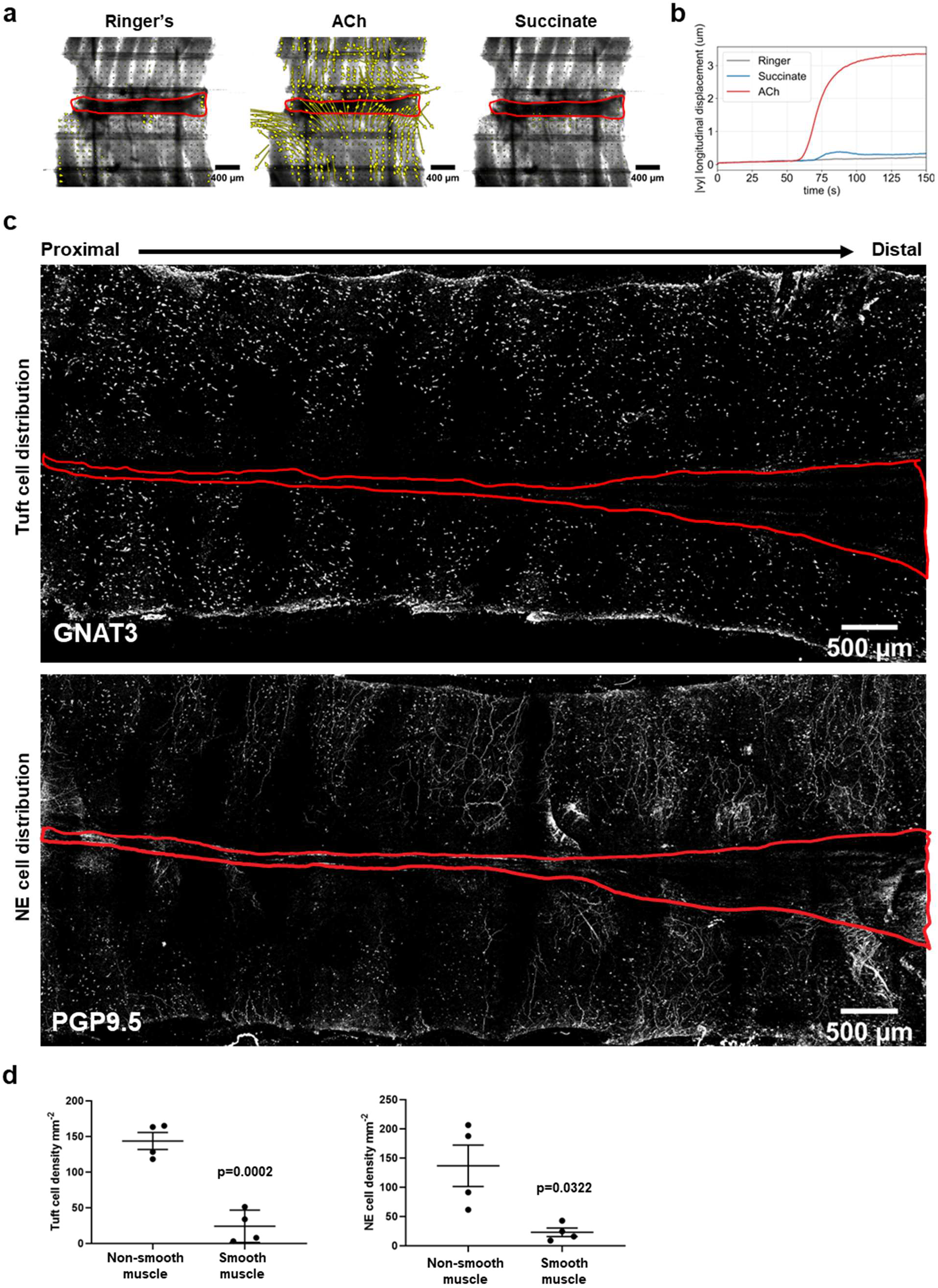
Sentinel cells are sparsely distributed over smooth muscle and succinate fails to elicit smooth muscle contraction. (a) Optical-flow vector fields at the time of peak displacement under three conditions: Ringer’s solution (negative control), 10 µM ACh, and 5 mM succinate. Yellow arrows indicate the cumulative tissue displacement from the start of the recording to the peak-response frame, computed by Farnebäck optical flow. With ACh addition, the vectors are long and spatially organized - directed downward in the upper tissue and upward in the lower tissue, converging on the posterior smooth muscle band of the trachea, consistent with vertical shortening driven by muscle contraction. Vectors are short and spatially incoherent following succinate application, reflecting the measurement noise floor. (b) Time course of the mean displacement magnitude in the y axis, |vy|, computed as the per-pixel absolute value of the vertical component of the optical-flow field. The ACh trace rises steeply from ∼60 s, indicating a robust smooth muscle contraction. The succinate trace fails to show robust smooth muscle contraction. Stimulus solutions were applied at t = 30s. (c) Immunofluorescence images of the whole-mount trachea. Top: tuft cell distribution visualized by GNAT3 staining. Bottom: NE cell distribution visualized by PGP9.5 staining. Red outlines demarcate the posterior membranous region of the trachea corresponding to the distribution of smooth muscle. Sentinel cell types are sparsely distributed in this region. (d) Quantification of tuft (left) and NE (right) cell density in the epithelium overlying smooth muscle versus non-smooth muscle regions. Both sentinel cell types show significantly reduced density over smooth muscle (Paired t-test).

**Extended Data Fig. 15.**
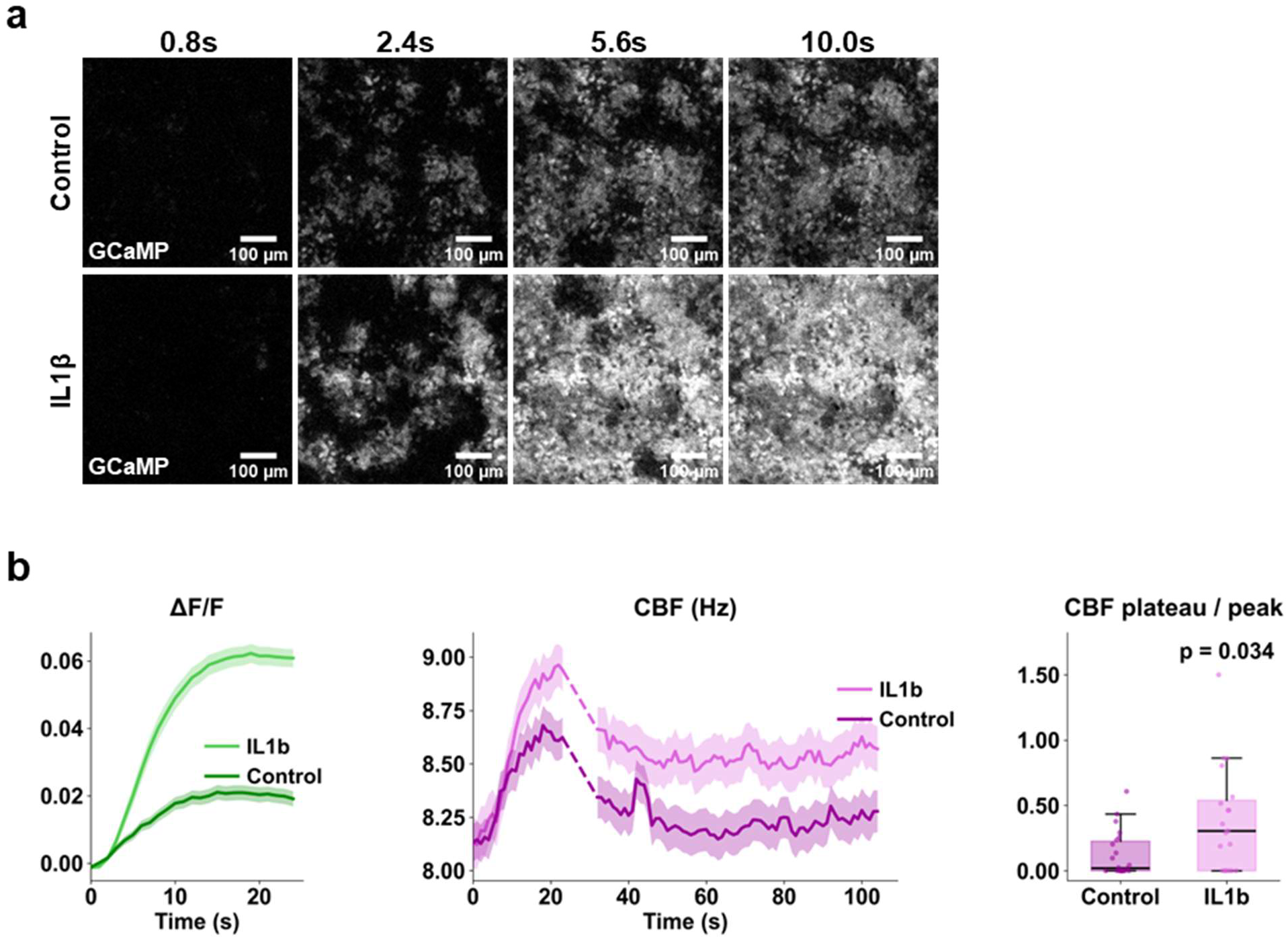
IL-1β enhances optogenetically induced DLDs and ciliary beat frequency (CBF). **(a)** IL-1β treatment enhances the intensity and domain size of optogenetically-induced DLDs in *AvilCreER; LV-FLEXon-mOPN4; LV-CBh-GCaMP8s* ALI cultures. Time-lapse imaging of calcium activation (GCaMP) shown for control (top) and IL-1β-treated (bottom) cultures. Time stamp in seconds shown on top. **(b)** Quantitative traces of calcium (ΔF/F, left) and CBF (Hz, middle) over time in control vs. IL-1β-treated cultures. Shaded bands denote SEM. IL-1β increases the peak amplitude of both calcium and CBF as well as the sustained plateau (average over 80-110s time window) of CBF. The dashed segment between 24–32s represents a time window in which CBF could not be calculated because the optogenetic blue light is switched off to minimize bleaching during this interval. Box plots (right) show the plateau-to-peak ratio for CBF, with significant elevation in IL-1β cultures (Mann-Whitney U test). Boxes show median and interquartile range; whiskers extend to 1.5×IQR.

**Extended Data Fig. 16.**
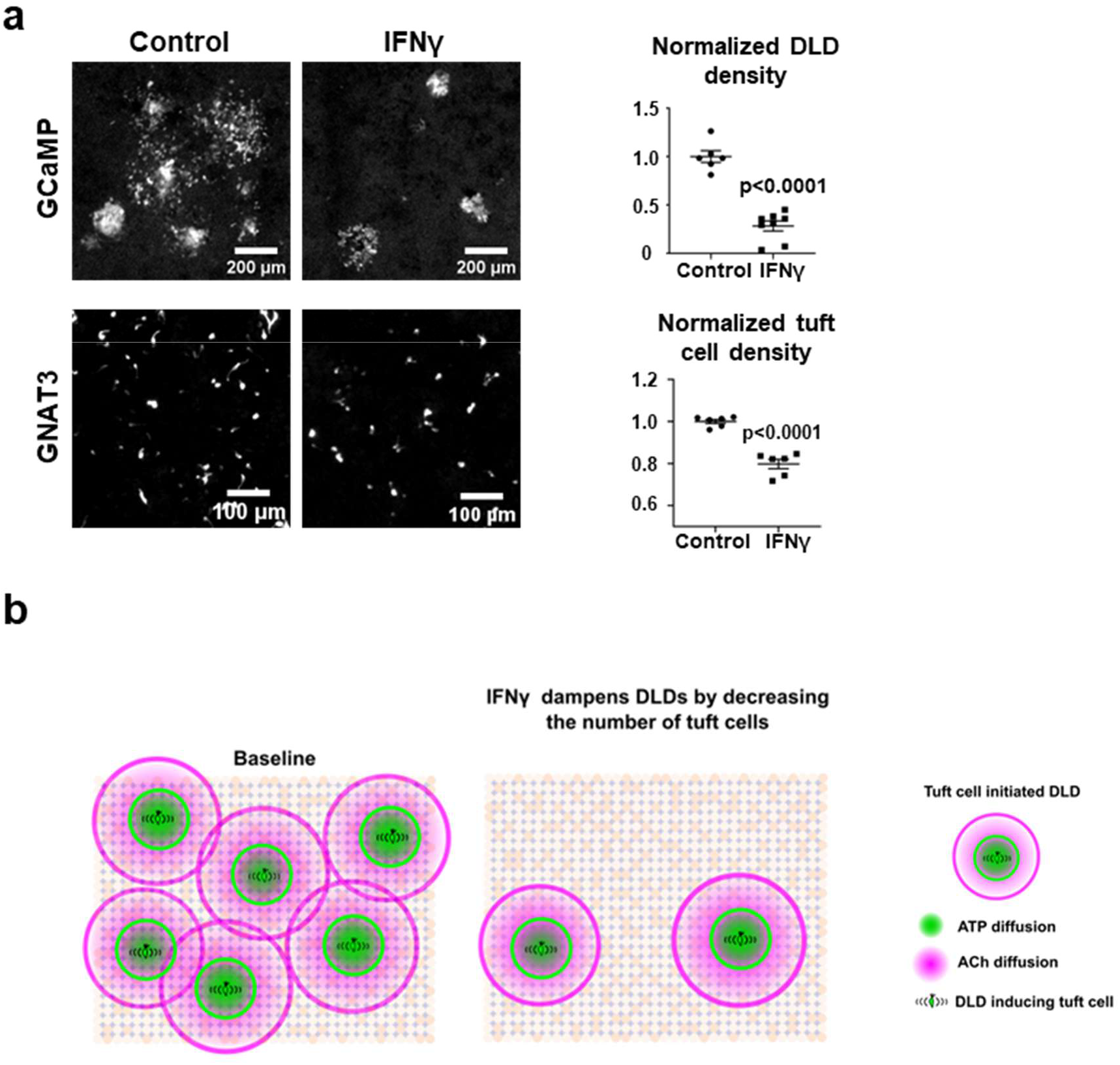
IFN-γ reduces DLD coverage through a reduction in tuft cell numbers. **(a)** IFN-γ results in fewer succinate-induced DLDs. Left and middle: Representative fluorescence images of ALI cultures from control and IFN-γ-treated preparations. Top row, GCaMP signal. Bottom row, GNAT3 immunostaining marking tuft cells. Right: Quantification of normalized DLD density (top) and normalized tuft cell density (bottom) (Welch’s t-test). (**b**) Schematic model of IFN-γ-mediated DLD dampening caused by decreases in tuft cell numbers.

**Extended Data Fig. 17.**
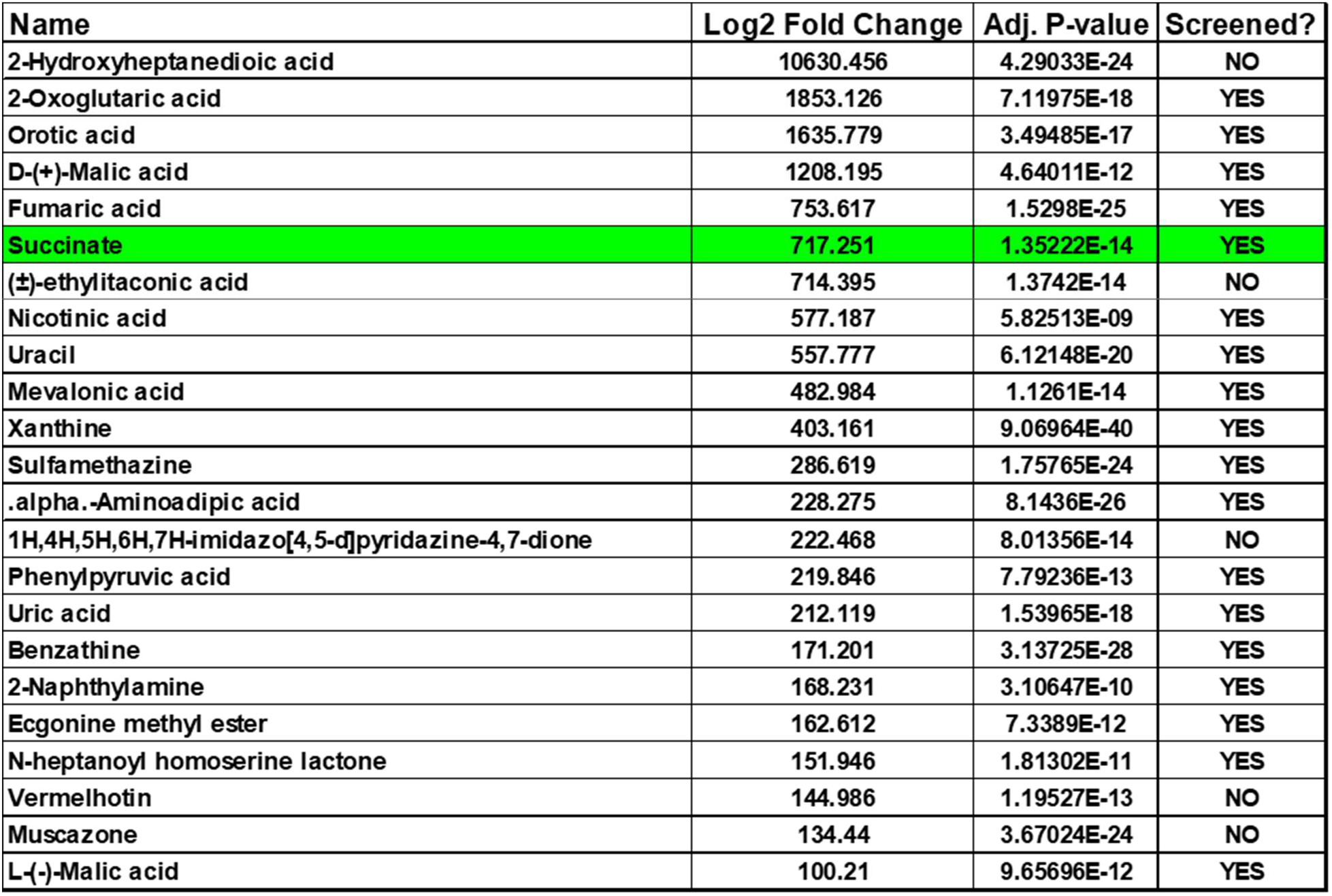
List of highly enriched metabolites in *Aspergillus fumigatus* conditioned media. List of metabolites identified by LC-HRMS as significantly enriched in *Aspergillus fumigatus*-conditioned media relative to control media, filtered for fold-change ≥ 100. Succinate (highlighted in green) was identified as the DLD-inducing metabolite.

## Materials and Methods

### Animals

Mouse lines *Nkx2.1Cre* ^51^ (JAX: 008661), *Nkx2.1CreER*^52^ (JAX: 014552), *LSL-GCaMP*^21^ (JAX: 031562), *LSL-Gq-DREADD*^30^ (JAX: 026220), *C57BL/6J*(JAX: 000664), *Ascl1CreER*^31^ (JAX: 012882), *AvilCreER*^28^ (JAX: 032027), *LSL-DTA*^29^ (JAX: 009669), *Chat^flox/flox^*^36^ (JAX: 016920), *Chat*-*GFP* ^44^ (JAX: 007902) were purchased from The Jackson Laboratory. *Scgb1a1CreER*^32^, *Foxj1Cre*^33^, *P63CreER*^34^, *Calhm1 KO*^38^ lines have been previously described. *Sucnr1 KO*^49^ mice were provided by the von Moltke lab at University of Washington. All mice were maintained on a 12-hour day-night cycle with *ad libitum* access to food and water in the Center for Comparative Medicine at the Massachusetts General Hospital, and all procedures were performed with Institutional Animal Care and Use Committee (IACUC)-approved protocols.

### Live calcium imaging with tracheal explants

To induce GCaMP expression in tracheal epithelial cells, adult *Nkx2.1CreER, LSL-GCaMP* mice received a single intraperitoneal injection of 200ul tamoxifen (20mg/ml, Sigma T5648) dissolved in corn oil (Sigma C8267). One week post-injection, mice were euthanized with carbon dioxide asphyxiation. Tracheas were dissected, cleared of connective tissue, and opened longitudinally along the ventral tracheal wall. Each trachea was then cut into four pieces with one cut along the trachealis muscle and the second cut along the middle tracheal cartilage ring. Explants were transferred to a laminar flow imaging chamber (Warner RC22 64-0228) and anchored to the glass coverslip bottom using a slice anchor (Warner 64-0246). Explants were continuously superfused (∼1ml/min) with a modified Ringer’s solution (125mM NaCl, 15mM NaHCO_3_, 2.4mM K_2_HPO_4_, 0.6mM KH_2_PO_4_, 1.2mM CaCl_2_, 1.2 mM MgCl_2_, 2.5mM D-(+)-Glucose, and 10mM HEPES, pH adjusted to 7.4 (NaOH)) at room temperature. 1 to 10mM succinate was dissolved in the modified Ringer solution to induce tuft cell-initiated ICWs. High K^+^ bath solution (50mM KCl, 75mM NaCl, 15mM NaHCO_3_, 2.4mM K_2_HPO_4_, 0.6mM KH_2_PO_4_, 1.2mM CaCl_2_, 1.2 mM MgCl_2_, 2.5mM D-(+)-Glucose, and 10mM HEPES, pH adjusted to 7.4 (NaOH)) was used to induce NE cell-initiated ICWs. Stimulus was applied using bath application that displaced the modified Ringer for 30 seconds to 90 seconds. Afterwards, modified Ringer was used to wash out the stimulus. Live calcium imaging was performed on a Nikon Eclipse 90i upright epifluorescence microscope equipped with a Prior Lumen 200 light source and Nikon DS-Qi1Mc CCD monochrome camera. The FITC filter set was used to excite GCaMP. Images were acquired with exposure time set between 100-500ms and imaging acquisition frequency between 2-5 Hz. Calcium image results were presented as ΔF/F GCaMP fluorescence where ΔF=F-F_baseline_. F_baseline_ is calculated by averaging over 5-10s GCaMP fluorescence prior to stimulus application.

### Air-liquid interface (ALI) culture

Tracheal basal stem cells were isolated from adult mice trachea using papain solution and plated on 804G-coated 6-well plates as described before^1^. Cells were expanded in the complete SAGM medium (Lonza, CC-3188) supplemented with TGF-β/BMP4/ WNT antagonist cocktails and 5 μM ROCK inhibitor Y-27632 (Selleckbio, S1049). Specific pre-seeding procedures based on genotype are outlined in Extended Data Fig. 1. To generate ALI culture, 150,000-250,000 basal stem cells were seeded onto trans wells on 24-well plates (Corning, 3470) with SAGM medium in both top and bottom chambers. One day post seeding, SAGM medium in both chambers was removed, and PneumaCult-ALI Medium (StemCell, 05001) was added only in the bottom chamber to create the air-liquid interface. ALI Medium was changed three times per week to allow for airway epithelium differentiation. 3 to 8-week-old ALI cultures were used for live calcium imaging. To induce CreER mediated recombination in ALI culture, 1 μM 4-Hydroxytamoxifen (4-OHT) (Sigma, H7904) was added into ALI medium for 5-7 days with the following exceptions: For ablation of tuft cells using *AvilCreER, LSL-DTA* line, 1 μM 4-OHT was added for 7-14 days. For ablation of NE cells using *Ascl1CreER, LSL-DTA* line, 1 μM 4-OHT was added for 9-14 days.

### Lentiviral transduction of basal stem cells

Lentiviral vectors were designed on VectorBuilder. Lentiviruses were packaged using standard lentiviral packaging protocol or directly purchased from VectorBuilder. 100,000; 200,000; and 300,000 basal stem cells were plated on 12-well plates one day prior to lentiviral transduction. On the day of transduction, wells with 40-60% confluence were selected to be transduced. Old medium was replaced with 1 mL fresh SAGM medium containing 5–17.5 µL lentivirus and 2 µg/mL polybrene. The next day, cells from each transduction were split onto 3 wells in 6-well plates to be expanded. Once reaching 50-80% confluence, cells underwent puromycin selection detailed below or further expansion to make ALI culture directly or FACS sorting. The following lentiviruses were used: *LV-Puro-CBh-GCaMP6s, LV-Puro-CBh-GCaMP8s, LV-CBh-GCaMP8s, LV-Puro-CBh-XCaMPR, LV-CBh-XCaMPR, LV-Puro-FLEXon-mOPN4, LV-Puro-CBh-ATP sensor, LV-CBh-ATP sensor, LV-Puro-CBh-ACh sensor*, and *LV-CBh-ACh sensor*.

### Puromycin selection with basal stem cells

Lentiviral transduced basal stem cells that contained a puromycin selection cassette were selected using decreasing concentration of puromycin for 3 days. On day 1, cells were resuspended and plated in SAGM with 3 µg/mL puromycin. On day 2, old medium was replaced with fresh SAGM containing 2 µg/mL puromycin. On day 3, old medium was replaced with fresh SAGM containing 1 µg/mL puromycin. Following selection, cells were expanded for ALI cultures.

### Live imaging with ALI cultures

Two methods were employed for live imaging with ALI cultures with consistent results between them. In the first method, ALI cultures were cut out from the trans wells and set up for imaging in the same way as tracheal explants described earlier. In the second method, ALI cultures were rinsed with modified Ringer to remove mucus buildup. 50ul of Ringer was then added on top of the trans wells to buffer the physical impact of adding stimulants to the top of the ALI culture.

The intact trans wells were then imaged. 100ul of stimulus solution including succinate, high K^+^, or DREADD agonist compound 21 (C21, biotechne, 6422) was added to the top chambers. 1-10mM succinate and 5-20 µM C21 were made in the modified Ringer solution. High K^+^ bath solution was the same as described for explants. Majority of the live imaging results were obtained with an Olympus IX81 inverted epifluorescence microscope. GCaMP/XCaMPR, ACh sensor/XCaMPR, and ATP sensor/XCaMPR dual-color live imaging were carried out on the IX81 or a Leica SP8 laser scanning confocal microscope. The framerate ranged from 0.2 - 2 Hz for image acquisition. High-speed high-resolution imaging of GCaMP, ACh sensor, and ATP sensor was carried out with a Nikon CSU Yokogawa Spinning Disk with framerate up to 16.7 Hz. Live imaging results were presented as ΔF/F fluorescence where ΔF=F-F_baseline_. F_baseline_ is calculated by averaging 15-30s GCaMP fluorescence prior to application of stimulants.

### Optogenetic activation of tuft cells in ALI cultures

ALI cultures were generated from *AvilCreER* basal stem cells transduced with *LV-Puro-FLEXon-mOPN4* and *LV-CBh-GCaMP8s* for optogenetic activation of tuft cells. To induce Cre-dependent *mOPN4* expression in tuft cells, the experimental group was treated with 1 μM 4-hydroxytamoxifen (4-OHT; Sigma, H7904) in the ALI medium for 4–7 days. One hour prior to imaging, ALI culture medium was replaced with modified Ringer media containing 1-5µM all-trans retinol (ATR) (Cayman Chemical, 18449) to provide the chromophore required for mOPN4 activation. Imaging was performed on an Olympus IX81 inverted microscope equipped with a FITC filter set; the ∼488 nm excitation light used for GCaMP8s imaging simultaneously activated mOPN4 in tuft cells, allowing concurrent optogenetic stimulation and calcium imaging.

For long-term optogenetic activation of tuft cells with EdU labeling, ALI cultures were generated from *AvilCreER* basal stem cells transduced with either *LV-Puro-FLEXon-mOPN4* and *LV-CBh-GCaMP8s* (for combined optogenetic activation and live calcium imaging) *or LV-Puro-FLEXon-mOPN4* alone (without calcium indicator). The *GCaMP8s*-expressing line was used for live calcium imaging to validate the protocol while the line without *GCaMP8s* was used for EdU quantification to avoid potential GCaMP8s-related phototoxicity during long-term illumination. Cre-dependent mOPN4 expression was induced as described above. Throughout the imaging period, ALI medium was supplemented with 2µM ATR to support mOPN4 activation and 1 µM EdU (Thermo Fisher Scientific, C10640) to label proliferating cells. Imaging was performed on a Nikon Eclipse Ti inverted microscope equipped with a FITC filter set; the ∼488 nm excitation light used for GCaMP8s imaging simultaneously activated mOPN4 in tuft cells, allowing concurrent optogenetic stimulation and calcium imaging for *GCaMP8s*-expressing ALI cultures and optogenetic stimulation alone for ALI cultures without *GCaMP8s*. Cultures were maintained in an environmental enclosure at 37 °C with CO₂ supplementation to maintain physiologic pH. After one day of imaging, the medium was replaced with fresh medium containing 2µM ATR and 1µM EdU, and cultures were rested for 1.5–2h without optogenetic activation before resuming imaging. At the end of the imaging period, cultures were fixed and EdU incorporation was detected by click chemistry per manufacturer’s instructions (Thermo Fisher Scientific, C10640). Cultures were subsequently immunostained with goat anti-SCGB1A1 (1:3000; Sigma, ABS1673) to mark secretory cells and rabbit anti-NGFR (1:100; Abcam, ab52987) to mark basal cells.

### Chemogenetic activation in ALI cultures

ALI cultures were generated from *AvilCreER; LSL-Gq-DREADD* basal stem cells for chemogenetic activation of tuft cells. 1μM 4-OHT was added into ALI medium for 5-7 days to induce Cre-dependent *Gq-DREADD* expression in tuft cells. 5-20 µM DREADD agonist compound 21 (C21, Bio-Techne, 6422) was used to activate Gq-DREADD.

For sparse chemogenetic activation of basal stem cells, secretory cells, and ciliated cells, ALI cultures were generated from *P63CreER, LSL-Gq-DREADD, Scgb1a1CreER, LSL-Gq-DREADD,* and *Foxj1Cre, LSL-Gq-DREADD* lines, respectively. Sparse labeling was achieved by titrating 4-OHT to low doses: 20nM for secretory cells and 10nM for basal stem cells. Since ciliated cell specification is already sparse in ALI cultures, Foxj1Cre driver was used without the need to add 4-OHT. 5µM C21 was used to activate sparsely labelled cells.

### High speed live imaging for simultaneous calcium and ciliary beat frequency (CBF) measurement

#### 1) CBF measurement

Two methods were used for CBF measurement at room temperature. For optogenetically induced DLDs, brightfield illumination was used to image cilia movement. For all other applications, 20 µg/ml Wheat Germ Agglutinin–CF488A (WGA-488; Biotium 29022) was used to label ciliated cells and image cilia movement. An Olympus DP23m camera and a Photometrics Moment camera mounted on an Olympus inverted IX81 microscope captured 40–50 frames per second, allowing CBF detection up to 20 Hz. CBF was computed by fast Fourier transform (FFT) over 4 s windows. For brightfield-based recordings, a notch filter was applied to remove the light-induced artifactual FFT peak near 10 Hz. CBF was calculated in 1 s steps with successive window centers, yielding a time-resolved CBF trace (Hz) for each analyzed unit. For air-liquid interface (ALI) cultures, CBF was calculated per individual ciliated cell, segmented using a temporal-activity-based watershed approach: a per-pixel temporal standard-deviation map was computed across the pre-stimulus segmentation window (exploiting the high-frequency intensity fluctuations of beating cilia), binarized by Otsu’s method with manual threshold refinement, morphologically refined (opening 2×2, closing 3×3, small-component removal <30 pixels²), and individuated by marker-controlled watershed seeded from local maxima of the smoothed Euclidean distance transform (Gaussian σ = 1 pixel, minimum seed spacing = 4 pixels). For explants, segmentation was omitted and CBF was calculated for the entire tissue. Change in CBF was expressed as a fractional change relative to a pre-stimulus baseline (ΔCBF/CBF), where the baseline was the mean CBF over a defined pre-stimulus window. Image processing used OpenCV and scikit-image.

#### 2 Calcium measurement

For brightfield illumination coupled with optogenetic induction of DLDs, blue light was applied for 20–30 s to enable (i) optogenetic activation of tuft cells expressing mOPN4 and (ii) visualization of DLDs through GCaMP fluorescence superimposed on the brightfield signal. For WGA-488-based detection of cilia movement, blue light was used throughout the imaging period. The combined green-channel signal contained WGA-488 (high-baseline live ciliated-cell label) and GCaMP (lower-baseline calcium indicator in all epithelial cells). Because the WGA-488 and brightfield signals were stable over the recording window, changes in green-channel pixel intensity were attributed to GCaMP fluorescence. Calcium responses were quantified as the relative change in green-channel intensity (ΔF/F), where F₀ was the mean signal in the pre-stimulus baseline window. DLDs were identified manually in ImageJ based on calcium activity. To correct for progressive photobleaching, a per-cell single-exponential decay F_bleach(t) = A·exp(−t/τ) + C was fit to each cell’s fluorescence trace using the signal-free portions of the recording (pre-stimulus baseline and post-response tail), excluding the stimulus response window. The fitted bleach curve was subtracted from the full trace prior to peak detection and downstream analysis. Curve fitting used nonlinear least-squares (scipy.optimize.curve_fit).

#### 3 Per-cell response quantification

For each ciliated cell within an identified DLD, the maximum calcium response (Max ΔF/F) was defined as the peak baseline-subtracted ΔF/F within the response window, and the maximum CBF response (Max ΔCBF/CBF) was defined as the peak fractional CBF change within the corresponding response window. Cells were matched between the calcium and CBF channels by DLD and cell identity.

#### 4 Per-DLD plateau to peak ratio

The plateau ratio was computed at the level of individual signaling DLDs rather than individual cells. For each wave, we first averaged across all cells in that wave at each timepoint, then computed the ratio of 30s late-window mean signal to the wave’s peak value. This wave-level aggregation matches the natural unit of biological independence in our data: cells within a wave respond to the same calcium signaling event originating from a single chemosensory cell in the same epithelial patch and are therefore not statistically independent observations. Treating each cell as independent would constitute pseudo-replication and inflate the apparent statistical power. Aggregating to wave level also stabilizes the ratio metric by averaging out per-cell denominator noise that would otherwise produce a heavy-tailed distribution of ratios. Comparisons between control and IL1β were performed using the two-sided Mann–Whitney U test on these per-wave ratios.

### Cytokine treatment of ALI culture

IL-1β (Peprotech 211-11B) and IFN-γ (Peprotech 315-05) were first dissolved in PBS as 500X to 1000X stock solutions and stored at -80°C. On the day of treatment, stock solutions were diluted in ALI culture medium to final concentrations of 20 µg/ml (IL-1β) and 50 µg/ml (IFN-γ). Mature ALI cultures (> 3-week old) were treated with IL-1β or IFN-γ for 5 consecutive days before live imaging.

### Assessment of tracheal smooth muscle contraction

Tracheal explants from *C57BL/6J* mice were used to assess smooth-muscle contraction. Tracheas were dissected from euthanized mice and placed in modified Ringer’s solution (described earlier). Surrounding connective tissue was removed under a dissecting microscope. Each trachea was cut open along the anterior midline and then divided into a proximal and a distal half. Distal halves contained a large area of smooth muscle near the carina and exhibited noticeable spontaneous contractile activity under our imaging conditions; they were therefore excluded from this study, and only proximal halves were used. Opened proximal explants were flat-mounted with the luminal side facing down in an imaging chamber (Warner RC-27L) on slice supporters that allowed solution flow both above and below the tissue. A slice anchor (Warner SHD-27LH/10) was placed on top to immobilize the explants. The chamber volume was maintained at 1 mL using a combined gravity-fed perfusion and peristaltic pump system, and tracheas were continuously superfused with modified Ringer’s solution at ∼1 mL/min at room temperature. For stimulation, 5 mM succinate (to induce tuft cell-initiated DLDs) or 10 µM acetylcholine (ACh; positive control for smooth-muscle contraction) was dissolved in modified Ringer’s solution and applied by gravity-based bath exchange, displacing the perfusing Ringer’s for 2 min. Tracheal explants were imaged on an Olympus IX81 inverted microscope with a 4× objective under brightfield illumination, and images were acquired at 5 frames/s.

Raw image stacks were processed in Python using a custom pipeline. Briefly, time lapse movies were rigid-body registered to the first frame using pystackreg to correct for chamber drift. Following registration, the void border introduced by translation/rotation alignment was identified per stack using a temporal-variance criterion (pixels with zero variance across all frames were classified as out-of-field), and the stack was cropped to the common valid rectangle. A tissue mask was generated by thresholding the 8-bit reference frame using Otsu’s method on the inverted image. Tissue motion was quantified using baseline-anchored Farnebäck optical flow as implemented in OpenCV (cv2.calcOpticalFlowFarneback). Flow was computed between the first frame (frame 0) and every subsequent frame *t*, yielding cumulative displacement vectors at every pixel. Farnebäck parameters were: pyr_scale = 0.5, levels = 4, winsize = 35, iterations = 5, poly_n = 7, poly_sigma = 1.5. The pixel-wise flow field (vx, vy) was used to compute several per-frame summary metrics. Tracheas were mounted with the smooth-muscle band horizontal in the image, so that muscle contraction shortens the tissue vertically (along the y-axis of the image). The headline contraction metric was therefore |vy|, the per-pixel absolute value of the y-component of the optical-flow field, averaged over the tissue mask.

### Immunostaining of ALI cultures and tracheal explants

ALI cultures and tracheal explants were washed with PBS and fixed in 4% PFA in PBS for 12-15min at room temperature. Fixed samples were rinsed in PBS twice before incubation with primary antibodies diluted in antibody buffer containing PBS + 0.3% Triton X-100 + 1% BSA overnight at 4°C. After washing 3 times in PBS, each time for 5min, samples were incubated with secondary antibodies containing DAPI (1:5,000) for 1-2 hours at room temperature. After that, samples were washed 3 times in PBS, each time for 5min, and then mounted with Fluoromount-G (SouthernBiotech, 0100-01). The following primary antibodies were used: rabbit anti-PGP9.5 (1:500; 14730-1-AP, Proteintech); goat anti-SCGB1A1 (1:3000; Sigma, ABS1673); mouse anti-Acetylated Tubulin (ATUB) (1:1000; Sigma, T6793); rabbit anti-NGFR (1:100; Abcam, ab52987); chicken anti-GFP (1:2000; Abcam, ab13970); mouse anti-TUJ1 (1:1000; Biolegend, 801202); mouse anti-HA (1:500; Thermo Fisher Scientific, 26183). All secondary antibodies were Alexa Fluor conjugates (488, 555, and 647) and used at 1:500 – 1:1000 dilution (Thermo Fisher Scientific). Images were obtained using Olympus Fluoview FV10i confocal microscope, Olympus IX81 epifluorescence microscope, and Leica SP8 laser scanning confocal microscope. Tuft and NE Cell numbers were counted based on immunostaining of markers for each of the respective cell types using ImageJ imaging software.

### Metabolites extraction

To selectively isolate the small-molecule fraction, CM from *Aspergillus fumigatus* cultures and uninoculated control media samples were first fractionated using a 3 kDa molecular weight cutoff (MWCO) (Sigma, UFC500396) filter via centrifugal filtration. An aliquot of 500 µL of the resulting filtrate (<3 kDa) was transferred to a 5.0 mL Eppendorf tube and mixed with 1000 µL of a methanol:chloroform mixture (3:1, v/v) and 20 µL of internal standard (anhydrous L-leucine-enkephalin acetate; 5 mmol/L). The mixture was vortexed briefly, incubated at -20°C for 5 min, and centrifuged at 10,000 rpm at 4°C for 10 min to induce biphasic partitioning. The upper polar phase (1 mL) was transferred to a clean tube and dried down completely using a SpeedVac vacuum concentrator (Thermo Scientific) at 35°C. The dried extracts were reconstituted with 100 µL of acetonitrile:water (70:30, v/v), vortex-mixed, and transferred into autosampler vials equipped with micro-inserts for LC–HRMS analysis. Reagent blanks were prepared by replacing the CM sample with 500 μL of Milli-Q water and subjecting it to the identical extraction steps. A pooled quality control (QC) sample was prepared by aliquoting 20 μL of each reconstituted study sample. The QC sample was both utilized to drive iterative intelligent data acquisition (AcquireX workflow) for comprehensive MS2 fragmentation and injected at regular intervals throughout the analytical sequence to monitor instrumental stability, retention time drift, and reproducibility.

### Liquid chromatography–high resolution mass spectrometry (LC-HRMS)

Metabolite extracts were analyzed by untargeted LC-HRMS using a Dionex Ultimate 3000 LC system consisting of a binary pump, an autosampler, and a column compartment, coupled to an Orbitrap ID-X Tribrid Mass Spectrometer (Thermo Scientific). Chromatographic separation was achieved in hydrophilic interaction liquid chromatography (HILIC) mode using a SeQuant® ZIC®-pHILIC column (150 mm × 4.6 mm, 5 µm particle size; Merck Millipore) at a flow rate of 300 μL/min. The total run time was 25 min using a multi-step mobile phase gradient. Mobile phase A consisted of 20 mM ammonium carbonate with 0.1% ammonium hydroxide in water, and mobile phase B was 100% acetonitrile. The elution gradient was programmed as follows: 0–2 min, 80% B; 2–18 min, linear gradient to 5% B; 18–20 min, hold at 5% B; 20–20.1 min, return to 80% B; and 20.1–25 min, re-equilibration at 80% B. The HILIC column and autosampler compartments were maintained at 25°C and 4°C, respectively. An injection volume of 5 µL was used for all samples. Data were acquired in centroid mode using a heated electrospray ionization (H-ESI) interface operating in both positive and negative polarities via separate analytical runs. The ionization parameters were optimized as follows: sheath gas flow rate, 30 arb; aux gas flow rate, 5 arb; sweep gas flow rate, 0 arb; spray voltage, 4.0 kV (positive mode) and 3.2 kV (negative mode); capillary temperature, 320°C; S-lens RF level, 50.0%; and aux gas heater temperature, 150°C. Full MS scan parameters were set to: resolution, 70,000 (at m/z 200); automatic gain control (AGC) target, (3x10^6^); maximum injection time (IT), 200 ms; scan range, m/z 70–1200. Data-dependent MS2 (ddMS2) selection was governed dynamically by the AcquireX™ intelligent data acquisition workflow (Thermo Scientific) to drive deep structural fragmentation across repetitive injections. The ddMS2 parameters were set to: resolution, 17,500; AGC target, (2x10^5^); maximum IT, 100 ms; isolation window, 0.4 m/z; stepped normalized collision energy (NCE), 20%, 30%, and 40%; minimum AGC target, 2x10^3^; and dynamic exclusion duration, 10 s. Real-time internal mass calibration was performed automatically using the EASY-IC™ system. Instrument control and data acquisition were managed via Xcalibur™ Software (Thermo Scientific), seamlessly paired with the AcquireX™ intelligent data acquisition workflow to generate automated background ion exclusion lists from blank injections alongside inclusion lists from pooled QC injections.

### LC-HRMS Data Processing and Small Molecule Identification

Untargeted metabolomic LC-HRMS raw data files were processed using Compound Discoverer software (version 3.3, Thermo Scientific) utilizing a standardized workflow template. Chromatographic peak detection, alignment, and feature extraction were performed using the Detect Unknown Compounds and Align Retention Times nodes, applying a mass tolerance of 5 ppm and a maximum retention time shift of 0.2 min. The minimum peak intensity threshold for feature extraction was set to 1x104 counts. Metabolite identification followed a multi-tiered orthogonal scoring approach. Structural formulas were predicted via the Predict Compositions node based on exact monoisotopic mass, isotope pattern spacing, and intensity distributions. Putative identifications were assigned via the Search Mass Lists and Search ChemSpider nodes across the HMDB, KEGG, and PubChem databases. High-confidence structural annotations (MS2 level) were achieved via the Search mzCloud node, requiring a minimum fragmentation library forward/reverse match score of 70. Definitive structural validation of key diagnostic metabolites, including succinate, was confirmed by matching retention times and accurate tandem mass spectra (MS/MS) fragments with authentic chemical reference standards. Prior to statistical assessment, peak areas were normalized using Quality Control (QC) Normalization via Local Polynomial Regression (LOESS) to correct for analytical variance across the injection sequence. Log2 fold-changes and statistical significance were calculated using the Differential Analysis node based on a pairwise Student’s t-test across designated study groups. Volcano plots were generated within the software by plotting the negative log10 of the p-value on the y-axis against the log2 fold-change on the x-axis. Thresholds for significantly altered features were strictly set at an absolute log2 fold-change ≥ 1.0 and a p-value <0.001.

### LC-HRMS Data Availability

The untargeted LC-HRMS raw data files, data-dependent MS2 (ddMS2) fragmentation spectra, processed peak lists, and chemical feature metadata generated during this study have been deposited in the MetaboLights repository under the accession number MTBLS14631.

### Quantification of succinate and benzathine concentrations in *Aspergillus* CM using LC-HRMS

For the quantitative determination of succinate and benzathine, two independent calibration curves were generated using authentic standards for each analyte. Standards were prepared in duplicate across a concentration range of 0.25 to 400 µM. Calibration curves were constructed by plotting analyte peak areas against nominal concentration, and linear regression was applied to establish the response relationship. These calibration models were used to quantify analyte levels in extracted conditioned media (CM) from Aspergillus fumigatus cultures. Quantification was performed by interpolation of sample responses within the corresponding calibration range.

### Statistical analysis

Statistical analyses were carried out with GraphPad Prism software and Python statistical packages including statsmodels, scipy.stats, and scipy.optimize. Specific statistical tests are detailed in the figure legends and relevant methods sections. A p value of less than 0.05 was considered significant.

### Human Ethics and Consent to Participate declarations

Not applicable

## Acknowledgments

We thank the members of the Rajagopal lab, Chivukula lab, and Charles Lin for constructive criticism. We thank Sam Wattrus for access to the spinning disc confocal microscope. We thank the Center for Comparative Medicine at the Massachusetts General Hospital for the maintenance of our mouse colonies. We thank Maris Handley and the MGH Simches Flow Core for FACS services. JX is a New York Stem Cell Foundation – Druckenmiller Fellow. JR would like to thank the Kayden family for Bernard and Mildred Kayden MGH Research Institute Chair; the Maroni family for the Kevin and Maroni MGH Research Scholar Award; and the Ro and Joanne Fallon Family for their donation to the laboratory.

## Funding

New York Stem Cell Foundation Grant 2023A000076 (JX); Harvard College Research Program (MH); Herchel Smith-Harvard Research Fellow (MH); National Institutes of Health R01CA279173, R35GM158046 (RM); National Institutes of Health R01HL148351 (JR); Howard Hughes Medical Institute Faculty Scholar (JR).

## Author contributions

Conceptualization: JX, JR. Methodology: JX, RVO, EJ, JLR, KNJ, MKM, RM, JMV, JR. Investigation: JX, MH, RVO, EJ, JLR, KNJ, HZ. Funding acquisition: JX, MH, PM, MKM, RM, JMV, JR. Writing: JX, MH, RVO, JR.

## Competing interests

Authors declare that they have no competing interests.

## Data and materials availability

The metabolomics data have been deposited to MetaboLights repository with the study identifier MTBLS14631. All other data are available in the main text or the supplementary materials.

## Notes

### Competing Interest Statement

The authors have declared no competing interest.

