## Supplementary Video Legends for "Rare Epithelial Sentinel Cells Establish Functional Domains of the Airway"

**Supplementary Video 1. Succinate induces Intercellular calcium waves (ICWs) in ALI culture.** Bath application of 10mM Succinate induces multiple ICWs in ALI culture. These ICWs originate from discrete foci distributed throughout the epithelium and propagate to eventually cover the entire tissue.

**Supplementary Video 2. Chemogenetic activation of tuft cells induces robust ICWs.** Chemogenetic actuator *GqDREADD* is expressed in tuft cells using *AvilCreER* driver. Application of DREADD ligand C21 induces robust ICWs in ALI culture with *GqDREADD* expression. No ICWs are induced in control ALIs that do not express *GqDREADD*.

**Supplementary Video 3. Optogenetic activation of tuft cells induces robust ICWs.** Optogenetic actuator mOPN4 is expressed in tuft cells using *AvilCreER* driver. Blue light is used to both activate mOPN4 and image GCaMP-based calcium signaling. Robust ICWs are induced in ALI culture with *mOPN4* expression. No ICWs are induced in control ALIs that do not express *mOPN4*. A few patches of immediate calcium activation are observed in control ALIs due to leaky expression of *mOPN4* in non-tuft cells.

**Supplementary Video 4. High $K^+$ induces ICWs in ALI culture.** Bath application of 50mM $K^+$ induces multiple ICWs in ALI culture. These ICWs originate from discrete foci distributed throughout the epithelium and propagate to cover large fractions of the entire tissue.

**Supplementary Video 5. ACh diffusion delimits tuft cell-initiated ICW domain size.** A dual ACh/$Ca^{2+}$ sensor reveals that the $Ca^{2+}$ wave (XCaMPR signal) slightly lags behind the diffusion of ACh (ACh signal) following succinate application.

**Supplementary Video 6. ICWs propagate across physical cuts.** An ICW propagate across physical cut created by a tungsten needle, indicating that direct cellular continuity is not required for ICW transmission.

**Supplementary Video 7. Tuft cells generate three types of DLDs.** A dual ATP/$Ca^{2+}$ sensor reveals three types of tuft cell-initiated DLDs following succinate application. Type A: A small ATP wave (green) surrounded by a larger $Ca^{2+}$ wave (magenta). Type B: A large $Ca^{2+}$ wave with no detectable ATP wave. Type C: A small $Ca^{2+}$ wave perfectly co-localized with an ATP wave.

**Supplementary Video 8. Tuft cells secrete ACh from the basolateral membrane.** High-speed, high-resolution spinning-disk confocal microscopy of ACh reporter reveals that ACh sensors are activated specifically at the basolateral membrane following succinate application.

**Supplementary Video 9. Tuft cells secrete ATP from the basolateral membrane.** High-speed, high-resolution spinning-disk confocal microscopy of ATP reporter reveals that ATP sensors are activated specifically at the basolateral membrane following succinate application.

**Supplementary Video 10. Local 1.5M KCl administration evokes spatially restricted DLDs.** Local administration of 1.5M KCl with a green fluorescent tracer evokes spatially restricted DLDs, confined to the area of tracer diffusion (grey haze).

**Supplementary Video 11. Succinate fails to induce robust smooth muscle contraction.** Brightfield time-lapse microscopy with tracheal explants demonstrates that succinate fails to induce robust smooth muscle contraction. Ringer’s solution is used as negative control whereas ACh is used as positive control which induces robust smooth muscle contraction.
